# Shaping the active site from a distance: Unravelling the regulatory role of the conserved N-terminal β-flap of thiol peroxidases through structure-function characterization of the Staphylococcal ortholog

**DOI:** 10.64898/2025.12.11.693838

**Authors:** Manjari Shukla, Sushobhan Maji, Amit Kumar Das, Amit Mishra, Sudipta Bhattacharyya

## Abstract

Bacterial thiol peroxidases (Tpxs) are members of atypical two-cysteine peroxiredoxin, which detoxify hydroperoxides during host-pathogen interactions. Despite extensive research on bacterial Tpxs, the precise structure-based functional characterization of Staphylococcal Tpx (SaTpx) remains elusive. Herein, we cloned and purified SaTpx along with its plausible physiological redox partners to reconstitute its electron relay system. We solved the high-resolution crystal structure of SaTpx, revealing the essential active-site catalytic triad (Cys 60, Thr 57, Arg 128). The crystal structure of SaTpx reveals the presence of the highly conserved N-terminal β-flap, absent in its mammalian orthologs (Peroxiredoxin 5). To investigate the plausible role of this highly conserved β-flap, we cloned and characterized a deletion mutant, NΔ15-SaTpx, lacking this region. NΔ15-SaTpx showed reduced ability to mitigate organic hydroperoxides *in vitro*; partially impaired to prevent ROS-mediated ds-DNA nicking and failed to protect host *E. coli* cells from exogenous ROS stress. The ultra-high-resolution crystal structures of NΔ15-SaTpx and NΔ15-SaTpx with substrate mimic revealed a distorted catalytic triad (with flipped Arg 128), explaining its partially impaired catalytic activity. This study, for the first time, elucidates the critical regulatory role of the N-terminal β-flap in stabilizing the catalytic triad of bacterial Tpx, providing insights for future inhibitor designing.

**Highlights:**

- Ultra high-resolution crystal structures of SaTpx and NΔ15-SaTpx
- Ultra-high resolution crystal structure of hydroperoxide substrate mimic bound NΔ15-SaTpx
- β-flap deletion mediated distortion of catalytic triad in NΔ15-SaTpx
- The novel regulatory role of the conserved N-terminal β-flap of Thiol peroxidases.

## 1. Introduction

Redox reactions are ubiquitous to any life form that thrive on earth, therefore, the enzymes and protein cofactors which are crucial for cellular redox homeostasis play indispensable roles in central life processes (1) which may range from shuttling of electrons through the electron transport chain during oxidative respiration to bombard noxious reactive oxygen/nitrogen species to the invading pathogens during innate immune defence (2–4). Especially in aerobic organisms, cellular redox homeostasis plays the pivotal role in their survival as the deleterious forms of molecular oxygen and its downstream reaction products must be mitigated in real time in order to stop the oxidation of the essential biological macromolecules (5). In this context, thiol disulfide based redox systems play the most trivial role to keep the deleterious reactive oxygen/nitrogen species at bay (1). Hence the enzymes and electron carrier proteins of thiol disulfide redox pathways are ubiquitous and essential, especially for aerobic life forms. Intriguingly, in intracellular pathogenic bacteria, thiol disulfide redox pathway proteins are also found to be important for the successful colonization and onset of diseases (6, 7). In *Mycobacterium tuberculosis* and *Staphylococcus aureus* thiol disulfide redox pathway enzymes and/or proteins are involved in the antioxidative defence mechanism against the host innate immune system generated reactive oxygen/nitrogen/sulphur species (8, 9). Inactivation of the key players of these pathways, hence found to cripple the intracellular bacteria to pursue a successful infection event. In this context, bacterial peroxiredoxins (Prxs) are the most robust frontrunners of thiol disulfide based redox defence system as they can directly neutralize incoming toxic organic hydroperoxides and lipid hydroperoxides to corresponding non-toxic alcohols with the aid of electrons shuttled successively through the cognate thioredoxins (Trxs) and thioredoxin reductases (TrxRs) (or AhpF) (10, 11). In these reactions, NADPH, the labile redox cofactor of thioredoxin reductase (or AhpF) is used as the terminal electron doner. The electrons are transferred from NADPH and shuttled by a chain of proteins (TrxR to Trx to Prx or AhpF to AhpC) through successive thiol disulfide formation/exchange/resolution reactions to ultimately reduce organic hydroperoxides to their corresponding alcohols (11).

Importantly peroxiredoxins are the broader subfamily having common ancestor which contains thioredoxin fold (12) Among all Trx superfamily, peroxiredoxins contain a highly conserved peroxidatic cysteine at N terminus (13). Further, based on the sequence analysis, Prx is classified into six distinct subfamilies. These six subfamilies; Prx1 (AhpC/Prx1), Prx6, Prx5, Tpx and BCP/PrxQ and AhpE are unambiguously classified based on distinct sequence and structural features (14). These subfamilies vary in oligomeric states, interfaces between two monomers and the location of resolving cysteine.

Additionally, Prxs have mechanistic classification which includes 1 cys, typical 2 cys and atypical 2 cys peroxiredoxin (15). In the classification 1 cysteine prx has only 1 cysteine and during catalysis regeneration of reduced state is accomplished by non-cysteine cellular reductants like glutathione and ascorbic acid (16). The 2-cysteine peroxiredoxins, and as the name suggests, have two redox active cysteine residues. They use active site cysteine (C_P_) thiolate group (Cys-S^-^) to commence a nucleophilic attack to the incoming hydroperoxide substrate to convert it to non-toxic alcohol, while in this process, the attacking thiol group is converted into cysteine sulfenic acid (Cys-SOH). In order to prevent further irreversible oxidation of the cysteine sulfenic acid to cysteine sulfinic acid (Cys-SO_2_H), as well as to restore the initial thiolate form of the peroxidative cysteine (C_P_), the thiol group (-SH) of resolving cysteine (C_R_) attacks the Cys-SOH to form a covalent disulfide bond. In later stages of the reaction cycle, the disulfide bond between the peroxidative (C_P_) and resolving (C_R_) cysteines of Prx is reduced by electrons received from reduced thioredoxin (Trx) (11). This is the mechanism of reduction of disulfide bond in most of the 2 cysteine peroxiredoxins with few exceptions. Majorly, both peroxidative (C_P_) and resolving (C_R_) cysteine residues are present in the same protomer of Prxs, however, depending on the location of the resolving cysteine (C_R_), 2-cysteine Prxs have been divided into two major subclasses, 1. Typical 2-Cys peroxiredoxins and 2. Atypical 2-Cys peroxiredoxins (17). Two cysteine peroxiredoxin (typical/atypical) both have the conserved catalytic motif P-X-X-X-T/S-X-X-C with arginine further stabilizing the thiolate form of cysteine and participating in substrate binding and catalysis (18). Typical 2-Cys peroxiredoxins are so far been extensively studied from structural and functional aspects, as they play crucial roles in redox homeostasis and bacterial pathogenesis. In this class, the C_P_ and the C_R_ do not belong to the same protomer of the enzyme oligomer, and upon oxidation by the hydroperoxide substrate, an intermolecular disulfide bridge is formed. On the other hand, the relatively less characterized member of the family, is the atypical 2-Cys peroxiredoxins, where both C_P_ and C_R_ belong to the same protomer and upon substrate-mediated oxidation, an intramolecular disulfide bridge is formed (17). Atypical 2-cysteine peroxiredoxins are relatively less characterized compared to the other counterparts. However, emerging evidences suggest they also play critical roles in bacterial pathogenesis (19). Among the six subfamilies of peroxiredoxin enzymes, Tpx (exclusively in bacteria), Dot5 (in yeast; eukaryote), PrxQ (in plants) Prx5/PRDX5 (in animals), enzymatic mechanism of peroxide reduction is executed through the formation of intramolecular disulfide bond among C_P_ and C_R_ cysteine (12). For many bacterial peroxiredoxins, reduction of the catalytic disulfide is mediated by the dedicated flavoprotein reductase AhpF, whereas numerous other peroxiredoxin systems in both bacteria and eukaryotes are regenerated via the thioredoxin/thioredoxin reductase (Trx/TrxR) electron transfer pathway (20) (19).

The Tpx subfamily of bacterial 2-Cys peroxiredoxins, a well-studied Tpx in *E. coli*, (21) adopts an obligate homodimeric assembly with an A-type dimer interface. A distinctive N-terminal β-hairpin, found exclusively in bacterial Tpx proteins, contributes to a conserved hydrophobic collar around the active-site pocket and underlies their pronounced substrate preference for alkyl hydroperoxides.

Genomic analysis of *Staphylococcus aureus* reveals the presence of several thiol specific antioxidant enzymes such as tpx (SAOUHSC_01822), AhpC (SAOUHSC_00365), and bcp (USA300HOU_1858). There is another low molecular weight thiol redox system in many Gram-positive Firmicutes i.e Bacilliredoxin (Brx). Brx are thiol-disulfide oxidoreductases and utilize Bacillithiol to reduce protein mixed disulfides (S-bacillithiolated proteins), thereby restoring protein function following oxidative stress.

The global superbug like *Staphylococcus aureus*, which lacks the glutathione-based antioxidative defence systems, atypical 2 cysteine peroxiredoxin (tpx) has been found to be indispensable to combat the deleterious effects of host-generated reactive nitrogen species (22, 23). Moreover, multiple scientific evidences suggest, thioredoxin dependent antioxidative defence pathway of *Staphylococcus aureus* is crucial to establish successful infection events and sepsis (24), rendering the non-redundant proteins of this pathway as lucrative drug targets.

Earlier, one preliminary crystallization condition of WT-SaTpx was reported (25), however, no structure-function characterization of the protein was carried out. In the present work we set out to explore the structure based functional attributes of Staphylococcal atypical 2-Cys peroxiredoxin (SaTpx) which was found to receive electrons from its redox partner protein Staphylococcal Thioredoxin 1 (SaTrx1) and Staphylococcal Thioredoxin reductase (SaTrxR) using NADPH as the terminal electron donor. The ultra-high resolution crystal structures of the SaTpx in solo and its substrate mimic bound forms presented herein have been used to unravel the plausible catalytic mechanism of the enzyme. Moreover, the ultra-high resolution crystal structures fortified by detailed biochemical and cell-based analysis, for the first time, enabled us to unravel the novel regulatory role of the active site distant β-flap of prokaryotic atypical 2-cysteine peroxiredoxins to regulate its catalytic activity.

## 2. Material and Methods

### 2.1. Phylogenetic distribution analysis of thiol peroxidase (atypical 2 cysteine peroxiredoxin)

The amino acid sequence of Staphylococcal atypical two cysteine peroxiredoxin, which is also known as thiol peroxidase (SaTpx) is retrieved from KEGG organism database and also matched with NCBI proteins database by pairwise sequence alignment (26). *Staphylococcus aureus* MRSA252 which is the parental strain of *S. aureus* RN4220 was used as the reference organism to retrieve the amino acid sequence and the corresponding nucleotide sequence of Staphylococcal thiol peroxidase. The obtained SaTpx amino acid sequence was further used to find sequence homologs using the tool HMMER (27). From the HMMER results, few representative sequence homologs of SaTpx were picked, spanning various organisms representing all kingdoms of the three domains of life (bacteria, archaea, and eukarya). We have selected specifically the atypical 2 cysteine prx from different subfamilies of Prx i.e. Tpx, PrxQ, Prx5 and Bcp. Multiple sequence alignment was carried out using the picked amino acid sequences (by using Clausthal Omega (28)) and the alignment results were fed to iTol (29) (Interactive Tree of Life) online tool to perform phylogenetic analysis and to create the dendrogram showing the phylogenetic distribution of the picked amino acid sequences of thiol peroxidase homologs.

### 2.2. Multiple Sequence Alignment (MSA) of SaTpx homolog of atypical 2 cysteine peroxiredoxins with solved crystal structure/NMR structure

Furthermore, we also used NCBI-BLAST (30) tool to identify sequence homologs of SaTpx with previously solved crystal structures/NMR structures which has been submitted to PDB (SaTpx amino acid sequence was BLASTed against PDB database). Multiple sequence alignment of those SaTpx sequence homologs with known Crystal/NMR structures has been conducted and the secondary structural information of solved crystal structures (Crystal structure of *S. aureus* (PDB id: 3P7X) thiol peroxidase has been taken as a reference) has been overlayed on top of the multiple sequence alignment file (using ESpript 3.0 (31)) to analyse structural implications over amino acid sequence conservation. Once again three-dimensional structural coordinates of SaTpx homologs with known crystal/NMR structures were obtained from Protein Data Bank (PDB (32)). The monomeric and functional dimeric forms of these downloaded structures are superimposed on one another to identify the precise structural disposition of conserved catalytic site residues as well to find unique structural elements specific for bacterial and or eukaryotic thiol peroxidases. Some of the obtained structures also represent different oxidation states of the thiol peroxidase (reduced and oxidized). Structural superimposition of the oxidized and reduced conformers also allowed us to identify key structural discrepancies between the two redox states of the thiol peroxidase. PyMOL was used to visualize the three-dimensional structural information of the solved crystal/NMR structures of the SaTpx homologs. Likewise, PyMOL can be also used to generate structural superimposition of two or more three-dimensional structural coordinates to fetch important structural discrepancies between compared SaTpx homologs and/or their different redox conformers.

### 2.3. Cloning of SaTpx, deletion mutant NΔ15-SaTpx, R128A-SaTpx, SaTrx1 and SaTR in *E. coli* host

Genomic DNA was collected from the *Staphylococcus aureus* RN4220 strain (derived from NCTC 8325) overnight culture to use as a DNA template in PCR reactions. The *S. aureus* thiol peroxidase (SAOUHSC_01822), deletion mutant (NΔ15-SaTpx), thioredoxin (SAOUHSC_01100) and thioredoxin reductase (SAOUHSC_00785) open reading frames were amplified using the polymerase chain reaction. The forward and reverse primers were designed using the genomic DNA of *S. aureus* RN4220 strain. PCR amplification was followed by restriction digestion of ORFs, and plasmids with BamHI and SalI enzymes. The digested ORFs and plasmid were then subjected to 1% agarose gel for confirmation, and then the gel purified vector and clone were used for the ligation experiment at 37° and 16°C. The ligated clones were then transformed into cloning host *E. coli* DH5α, which were made chemically competent earlier with the CaCl_2_ method (33), and ligation was confirmed with colony PCR with gene specific forward as well as reverse primers followed by 1% agarose gel electrophoresis. pET28a vector was used for cloning of SaTpx, deletion mutant NΔ15-SaTpx while SaTrx1, SaTR were cloned in pQE30 vectors. Site-directed mutagenesis of SaTpx was performed at position 128 to obtain R128A-SaTpx mutant. Mutagenesis was performed using QuikChange II Site-Directed Mutagenesis Kit (Agilent), following the protocol. The recombinant plasmid DNA extracted from positive clones was then transformed into expression host BL21 (DE3) or M15 *E. coli* cells for overexpression of proteins.

### 2.4. Overexpression of recombinant proteins

We obtained heterologous expression of Staphylococcus aureus peroxiredoxin (SaTpx), its N-terminal β-flap deletion mutant (NΔ15-SaTpx), thioredoxin1 (SaTrx1) and thioredoxin reductase (SaTR) with *E. coli* expression host cells. SaTpx and NΔ15-SaTpx, R128A-SaTpx was expressed in the BL21(DE3) strain devoid of Lon and OmpT proteases to avoid protein degradation, whereas SaTrx1 and SaTR were expressed through pQE30 vector in *E. coli* M15 cells which also carries the pREP4 plasmid (kanamycin resistance) to allow lac repressor-mediated control. For overexpression and protein purification experiments, cryopreserved glycerol stocks of sequence confirmed clones were revived in LB broth supplemented with 50 µg/mL kanamycin (for pET28a-based expression constructs) and 50 µg/mL kanamycin and 100 µg/mL ampicillin (for pQE30 based expression constructs). Primary overnight cultures (incubated at 37°C with 200 rpm shaking) were diluted 1:100 in 2L LB medium under the same respective antibiotic selection pressure. Aerobic growth was maintained at 37°C, with 150 rpm agitation to mid-log phase (OD600 = 0.6), an optimal state for induction, where transcriptional activity was high. Expression of the recombinant protein was induced by adding the 100 µM IPTG. The post-induction cultures were incubated at 37°C for 4 hours for maximum production of soluble protein.

### 2.5. Purification of recombinant SaTpx, deletion mutant NΔ15-SaTpx, R128A-SaTpx and other associated proteins to homogeneity

The cells were harvested by centrifugation at 3,500 × g (at 4°C) for 40 minutes in a swing-bucket rotor. The collected pellet obtained after centrifugation was then dissolved in lysis buffer, also referred to as buffer A (10 mM Tris, 300 mM NaCl, 10 mM imidazole, pH 8.0). Sonication experiments were performed to lyse the cells at 4°C with 40 sec of heat and a 40 sec rest cycle in the probe sonicator. The lysed cells were then centrifuged at 13000 rpm for 80 minutes at 4 °C and the collected supernatant was applied to the Ni-NTA column, which was pre-equilibrated with equilibration buffer (Buffer A). The bound protein to the column was then eluted with buffer containing increasing concentrations of imidazole, like 10 mM, 20 mM, 50 mM, 75 mM, 100 mM, and 300 mM, and a constant concentration of 10 mM Tris and 300 mM NaCl adjusted to pH 8.0. The collected fractions were then checked for the most purified concentration of the desired protein with a 12% SDS PAGE experiment. The particular fraction was then concentrated with appropriate molecular weight protein concentrators and subjected to a pre-equilibrated size exclusion column (SEC) with 20 mM Tris and 50 mM NaCl concentration buffer adjusted to pH 8.0. The homogenous, highly purified protein fraction was collected, concentrated, and rapidly frozen at -80 °C with 5% glycerol.

### 2.6. Reconstitution of redox active disulfide-based electron relay system through coupled assay to determine the plausible *in vivo* electron acceptor of SaTpx

Two different sets of enzyme catalysis strategies have been performed to compare the catalytic efficiency of WT-SaTpx versus NΔ15-SaTpx. In the first reaction strategy, (termed as coupled assay) the fully functional thiol-disulfide redox enzyme cascade-based electron relay system has been constructed *in vitro*. This redox enzyme cascade involves Staphylococcal thiol-peroxidase (WT-SaTpx or NΔ15-SaTpx) and its upstream electron donor enzymes, Staphylococcal thioredoxin 1 (SaTrx1) and Staphylococcal thioredoxin reductase (SaTR). In this reaction cascade, the thiol peroxidase-mediated reduction of the incoming substrate, tert-butyl hydroperoxide (t-BOOH), results from the successive flow of electrons donated from the ultimate electron donor enzyme SaTR via the intermediate electron carrier, SaTrx1, through iterative thiol-disulfide exchange reactions. During this reaction, NADPH, the labile redox cofactor of SaTR is oxidized to NADP^+^. Hence, the catalytic reduction of tert-butyl hydroperoxide by the thiol-disulfide redox enzyme cascade can be spectrophotometrically monitored by the concomitant oxidation of NADPH to NADP+ by following the reduction of UV absorption at 340nm, where NADPH selectively absorbs. This reaction strategy was initially used to determine the most optimal hydroperoxide substrate of SaTpx **(Fig. S7a).**

Coupled assay entails the integration of thiol peroxidase activity with thioredoxin reductase and thioredoxin. This assay involves the NADPH as reducing agent (34). The reaction mixture contains the NADPH, SaTpx, SaTrx1, SaTR in the buffering system having 0.2 mM, 1.5 μM, 2 μM, 5 μM concentration respectively. The reaction gets started by adding the t-BOOH of 2 mM concentration in the reaction mixture and absorbance was measured quickly after t-BOOH addition at every 30 seconds and for time interval of 5min. Reaction mixture devoid of SaTpx was taken as control in the experiment. Coupled assay was also performed with mutant thiol peroxidase NΔ15-SaTpx and R128A-SaTpx mutant to evaluate the difference in catalytic efficiency of wild type and mutant form. The absorbances were measured using a UV-Vis spectrophotometer manufactured by Shimadzu called the UV-1900i.

### 2.7. Assessment of disulfide bond formation (free thiol group oxidation) in SaTpx as function of increasing t-BOOH concentration

In the second reaction strategy, a thiol peroxidase based catalytic conversion of hydroperoxide substrate has been considered in the absence of any other electron doner. The catalytic conversion of substrate hydroperoxide [tert-butyl hydroperoxide (t-BOOH) by the reduced form of thiol peroxidase (either WT-SaTpx or NΔ15-SaTpx) would also concomitantly generate the disulfide-linked oxidized form of the enzyme. In the absence of any electron donor, the formation of the disulfide-based oxidized form of the thiol peroxidase would be irreversible. The rate of formation of the oxidized thiol peroxidase hence would be directly proportional to the rate of catalytic conversion of substrate hydroperoxide to the corresponding alcohol and water (t-BOOH to t-BOH and H_2_O).

The Ellman’s reagent assay or the 5,5’-dithiobis-(2-nitrobenzoic acid) (DTNB) assay was employed to evaluate the presence of thiol groups in a sample (35). DTNB reacts with the thiol group to produce a yellow-coloured compound known as 5-thio-2 nitrobenzoic acid (TNB), which can be measured at 412 nm wavelength. The purpose of this experiment was to assess the oxidation of thiol groups present in the protein after the addition of t-BOOH in increasing concentrations (0μM, 2.5μM, 5μM, 7.5μM, 10μM, 25μM, and 50μM). SaTpx of final concentration 20 µM was added to the reaction mixture containing 20 mM Tris, 50 mM NaCl buffer. The reaction was initiated by adding t-BOOH and incubated for 1 minutes at 37 °C. The reaction was then stopped by adding 10% SDS. The DTNB of 500 µM final concentration was added and again, the reaction mixture was incubated for 30 min for colour development. The absorbance was measured at 412 nm. The experiment was also performed in the same manner with NΔ15-SaTpx and R128A-SaTpx mutant to compare the catalytic efficiency with wild type SaTpx.

### 2.8. Comparative analysis of Staphylococcal thiol peroxidase (SaTpx) catalytic activity against diverse hydroperoxide substrates

To determine which peroxide substrate SaTpx prefers, a coupled assay was conducted. This assay employs three distinct hydroperoxides: hydrogen peroxide (H_2_O_2_), tert-butyl hydroperoxide (t-BOOH), and cumene hydroperoxide. In each case, in addition to the hydroperoxide substrate, the reaction mixture contained NADPH, SaTR, SaTrx1 and SaTpx. For each reaction mixtures, the concentrations of NADPH, SaTR, SaTrx1, and SaTpx are as follows: 0.2 mM, 1.5 µM, 2 µM, and 5 µM, respectively, while the concentration of hydroperoxide (H_2_O_2_/t-BOOH/cumene hydroperoxide) was kept at 2 mM. The final pH of the reaction buffer, composed of 20 mM Tris and 50 mM NaCl, was 8.0. Initially, a suitable volume of SaTR, SaTrx1, or SaTpx was added and incubated at 4°C for one hour. Subsequently, the incubated protein mixture, NADPH, and an appropriate volume of 10x buffer were sequentially added to a microcentrifuge tube containing Milli-Q water. The reaction was initiated by the introduction of hydroperoxide substrates. The absorbance at 340nm was measured immediately following the addition of hydroperoxide, and subsequently repeated at 30 second intervals over the course of 5 min. As a control, a reaction mixture devoid of SaTpx was used for each instance. Absorbance at 340 nm was measured using a UV-Vis spectrophotometer. The K_m_ and V_max_ calculations have been done by taking varying concentrations of the substrate. The Origin software [OriginLab Corporation, Northampton, MA, USA.] used for fitting the data.

### 2.9. Catalytic activity comparison of wild type (SaTpx) and mutant staphylococcal thiol peroxidase (NΔ15-SaTpx)

The catalytic activity of SaTpx and NΔ15-SaTpx was also tested using DTNB, as well as a coupled assay with the same protocol and concentration used for SaTpx. K_m_ and V_m_ values were calculated as previously described. The secondary plot has also been plotted by taking three different concentrations of SaTrx1 (1 µM, 2 µm and 4 µM) to estimate true kinetic constants.

### 2.10. Biophysical assay to assess the three-dimensional structural change during oxidation of SaTpx/NΔ15-SaTpx

Circular dichroism spectroscopy was employed to detect the local unfolding of the SaTpx monomer as a function of cysteine oxidation in SaTpx/NΔ15-SaTpx upon exposure to the substrate tert-butyl hydroperoxide. This oxidation process triggers conformational changes, resulting in substantial modifications of the secondary structure of the protein. The experiment involved introducing varying concentrations of the oxidizing substrate t-BOOH into a reaction mixture containing 5μM SaTpx/NΔ15-SaTpx. Spectra ranging from 180 nm to 400 nm were recorded at a rate of 1nm/s. An online tool called BeStSel (36) was used to quantify the percentage changes in the alpha helix, beta turn, and loop contents of the protein structure.

### 2.11. *In vitro E. coli* cell protection assay with wild type and mutant Staphylococcal thiol peroxidase

The protective effect of the SaTpx/NΔ15-SaTpx clone in *E. coli* host cells under oxidative stress conditions was assessed using a modified version of the SPOT assay developed by Sahu S.R (37). *E. coli* cells containing an empty plasmid or the cloned construct (SaTpx/NΔ15-SaTpx) were grown with IPTG induction until they reached 0.4 OD at 600nm. After a 30-min incubation, oxidative stress was triggered with 5 mM, 10 mM and 15 mM tert-butyl hydroperoxide (t-BOOH). Following this, 5μl of various treated and serially diluted samples was spotted onto an agar plate. The plates were incubated at 37 °C for 16 h, after which the colonies were enumerated.

### 2.12. *In vitro* dsDNA protection assay from Reactive Oxygen Species (ROS) with wild type and mutant Staphylococcal thiol peroxidase

Peroxidases play a crucial role in preserving the structural integrity of dsDNA by counteracting a process called dsDNA nicking induced by reactive oxygen species. This protective function prevents breakage of individual strands within the dsDNA double helix, thus ensuring the stability of the genetic material. A dsDNA protection assay was performed as described previously with some modification (38) to assess the ability of SaTpx/NΔ15-SaTpx to protect dsDNA under oxidative stress. For the reaction mixture, 100 ng of pET28a plasmid DNA was mixed with 5 mM DTT, 15 μM FeCl_3_, and 20 μM SaTpx/NΔ15-SaTpx in 0.1 M HEPES-NaOH buffer at pH 7.0, and incubated at 37°C for 30 min. EDTA was added at a final concentration of 50 mM to terminate the reaction. A positive control was implemented by adding EDTA to the mixture prior to reaction initiation. SaTpx/NΔ15-SaTpx was excluded from the reaction mixture as a negative control. Ten microliters of each sample were loaded onto a 1% (w/v) agarose gel and electrophoresed at 100V for 45 min. The protected DNA content was calculated (using ImageJ software (39)) by comparing the intensities of the ethidium bromide-stained control and test sample DNA visualized under GelDock.

Additionally, TUNEL assay has been performed and TUNEL Assay Kit was purchased from Promega. To perform TUNEL analysis, bacterial samples were seeded into culture plates, and on the following day, cells were treated with different conditions as explained in the result section and after treatment, bacterial samples were used for TUNEL staining as per manufacturer’s instructions.

### 2.13. Crystallization of SaTpx, and NΔ15-SaTpx

Preliminary crystallization screening was performed on homogeneously purified target proteins using sparse matrix screening solutions (Crystal Screen I, Crystal Screen II, and Index Screen) sourced from Hampton Research. SaTpx & NΔ15-SaTpx crystallization was attempted with protein concentrations ranging from 10 mg/mL to 60 mg/mL using the sitting drop vapour diffusion method at ambient temperature and Hampton Research 96-well Linbro crystallization plates. After obtaining a limited number of initial crystallization results, a detailed screening of those crystallization conditions at room temperature was performed using the 60 mg/ml protein concentrations. A fine screening was conducted in which single diffraction quality crystals were grown at room temperature using hanging drop vapor diffusion method. In the SaTpx crystallization trial, 2 µl reservoir solution containing 1M Sodium citrate and 0.1M HEPES adjusted to pH 7.0 was mixed with 2µl of protein having a concentration of 60 mg/ml. While the NΔ15-SaTpx crystal has been grown with hanging drop vapor diffusion method in which crystallization drops were casted with 2µl 60mg/ml purified protein mixed with 2µl reservoir solution containing 1.8 M ammonium sulphate, 0.1 M HEPES pH 7.0 and PEG 400. The substrate mimic-soaked crystals with mutant NΔ15-SaTpx were obtained by soaking with buffer containing OtBH (O-tert-Butyl hydroxylamine hydrochloride) 6.6 mM or TBH 10 mM, and 1.8 M ammonium sulphate, 0.1 MHEPES pH 7.0 and PEG 400, for 5 min prior to flash freezing using liquid nitrogen.

### 2.14. Data collection and structure solution of SaTpx, NΔ15-SaTpx and substrate mimic OtBH/TBH soaked NΔ15-SaTpx

The crystals were picked under the stereomicroscope and soaked with respective buffers prior to data collection. Crystal loop picked single crystals were then flash frozen with liquid nitrogen. Since the crystallization mother liquor contains high amount of salt and/or PEG molecules, no additional cryo-preservatives were needed before flash freezing. The X-ray diffraction data collection was done at PX-BL21 beamline at RRCAT Indore India (37, 40). The diffraction images were collected from cryo-cooled crystals with 1° oscillation, using MarCCD 225 Rayonix detector. A full set of diffraction data was collected by rotating the mounted crystal, depending on the space group symmetry of the mounted crystal. After the full set of data collection, the diffraction images were indexed and integrated using XDS (41), followed by scaling in CCP4 suite (42), SCALA (43), while the Pointless (43) indicated the initial estimation of the space group. The scaling of SaTpx diffraction data was done in P2_1_2_1_2_1_, SaTpx with P2_1_2_1_2_1_ and NΔ15-SaTpx soaked with OtBH in P2_1_2_1_2_1_ space groups but for OtBH soaked mutant structure has to be reindexed scaled and refined in P2_1_ space group due to the presence of pseudo-hemihedral twining detected by “L-test”(44). Phasing was done with molecular replacement by using the either Phaser or Molrep (45) from CCP4 suite. The previously submitted crystal structure of SaTpx (PDB Id: 3P7X) served as the search model for molecular replacement-based phasing method. Afterwards, the iterative cycle of model building and crystallographic refinement (rigid and restrained) was done with COOT (46) and Refmac5 (47) (from CCP4 suite) respectively. The final cycles of crystallographic refinements were conducted through Phenix (48) suite. Furthermore, validation of the refined model was done with Molprobity (49) on Phenix suite. The parameters indicating data collection statistics crystallographic refinement, and quality of the final model are listed in **Table 2**.

### 2.15. Molecular Dynamics Simulation studies of SaTpx and NΔ15-SaTpx structure

Simulation analyses were conducted using the NAMD 2.14 software package (50). NAMD employs VMD (51) for preparation of input files. Ligand parameter and trajectory files were generated through Charmm-Gui Ligand Reader and Modeler (52). A solvation cube of 10Å was constructed using the isothermal-isobaric ensemble (NPT) in conjunction with Langevin dynamics. Multiple time stepping was applied with a time step of 1fs per integration cycle. The frequencies for saving trajectory (dcd), extended system (xst), restart frequencies, and output energies were all set to every 5000 steps. Minimisation was conducted for 1000 iterations prior to production run and simulation studies were performed for 100ns. The RMSF plot was created using Origin Software.

## 3. Results and Discussion

### 3.1. Atypical 2 cysteine peroxiredoxins orthologs are omnipresent in the three domains of life

The ubiquitous presence of atypical two-cysteine Prx homologs in three domains of life (bacteria, archaea and eukarya) **(Fig. 1a)** emphasises their functional indispensability in cellular redox homeostasis and redox signalling. From simple unicellular prokaryotes to complex multicellular eukaryotes, their functional variety and evolutionary preservation point out their high relevance in the context of redox control in biological systems. The unrooted phylogenetic tree further illustrates the occurrence of atypical 2-cysteine peroxiredoxin homologs in representatives from all major kingdoms of life, demonstrating their ubiquity and indispensable function throughout different life forms. All the members of this protein family share highly conserved catalytic motifs and well-preserved catalytic mechanism. In addition, the tree demonstrates the functional evolution of these enzymes, to suit the particular cellular requirements in different organisms. For example, higher plants and mammals have evolved multiple isoforms of peroxiredoxins, each targeted to a specific subcellular compartment, thus providing finely-tuned antioxidative protection. In higher eukaryotes peroxiredoxins are also involved in redox signalling. Prokaryotes, on the other hand, typically rely on fewer but highly efficient peroxiredoxins which protect them against oxidative stress, and so far, peroxiredoxin mediated redox signalling system has not encountered in prokaryotes.

**Figure 1:**
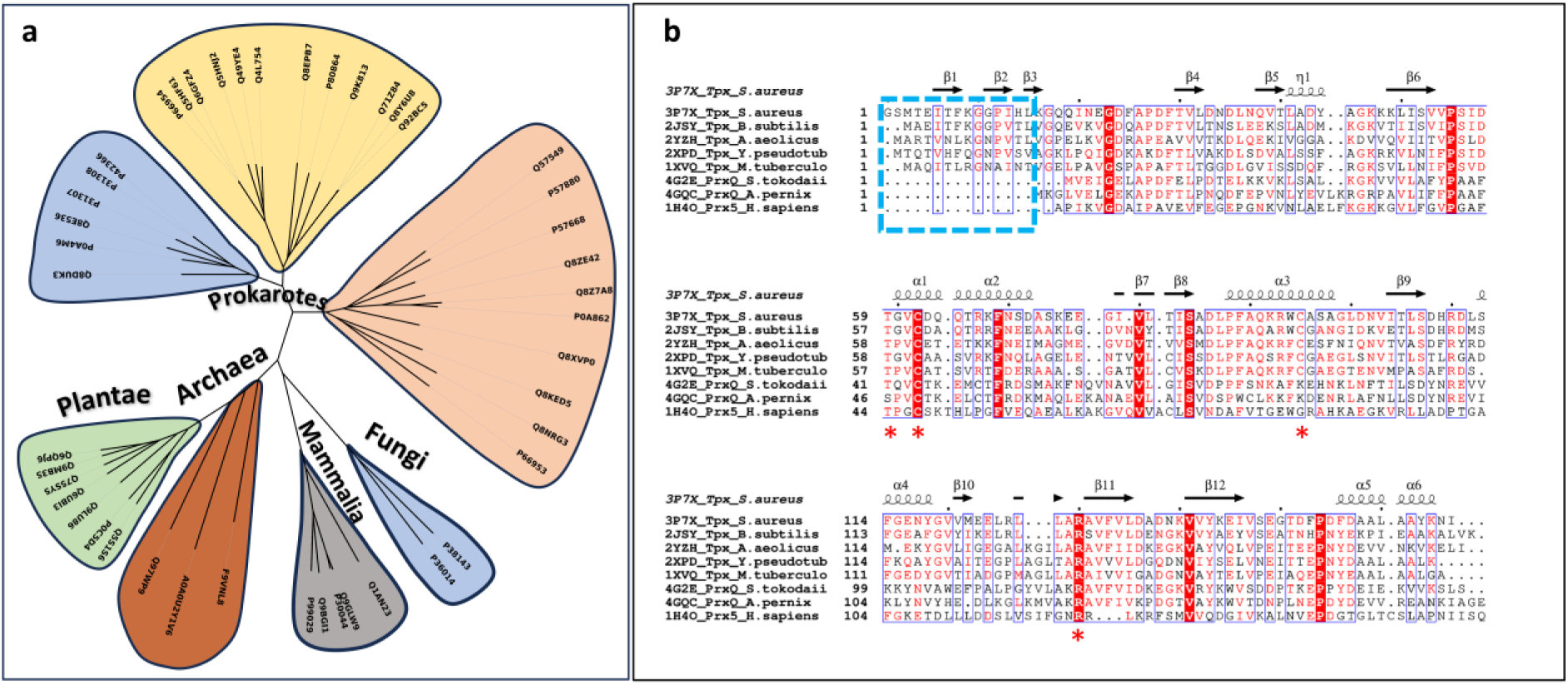
Presence of atypical two cysteine peroxiredoxin homologs across three domains of life. **1a:** The unrooted phylogenetic tree demonstrates the ubiquitous occurrence of atypical two-Cysteine Peroxiredoxin homologs throughout all major kingdoms of three domains of life with representatives identified in archaea, bacteria, protists, fungus, plants, and mammals. Each branch or cluster in the diagram represents a group of homologous peroxiredoxin sequences, with UniProt accession numbers indicating specific protein entries from various species. **1b: Multiple sequence alignment of atypical two-cysteine peroxiredoxins from diverse taxa having pre-determined crystal structure or NMR structure**. The sequence alignment is overlayed with known secondary structural information from bacterial atypical two-cysteine peroxiredoxin counterparts (also known as thiol peroxidases). The alignment reveals that prokaryotic (bacterial) atypical 2-cysteine peroxiredoxins possess a distinct N-terminal β-flap region, highlighted by the cyan box, which is absent in archaeal and eukaryotic homologs. Secondary structure elements are annotated above the alignment, illustrating the unique presence of this structural motif in bacterial sequences. The amino acid residues marked with red asterisks also demonstrates the highly conserved active site catalytic residues (except C_R_ whose position changes) of atypical two cysteine peroxiredoxins.

### 3.2. Multiple Sequence Alignment (MSA) of Staphylococcal atypical 2 cysteine peroxiredoxin (SaTpx) homologs with solved crystal /NMR structure uncovers the conserved catalytic triad and the unique N terminal β-flap of prokaryotic counterparts.

Amino acid-based multiple sequence alignment of SaTpx homologs with known crystal or NMR structures **(Fig. 1b)** reveals the high degree of conservation of amino acids constituting the active site catalytic triad (residues marked with an asterisk). Moreover, a distinct N-terminal β-flap region which is exclusively found in bacterial homologs was also identified (highlighted by the cyan box in **Fig. 1b).** Interestingly, this structural motif does not appear in peroxiredoxins from archaea, plants, or humans, indicating that it is a unique structural feature of prokaryotic enzymes. Importantly, human peroxiredoxin 5 (PRDX5-PDB id-1H4O (53)) represents the sole member of the atypical 2 Cys peroxiredoxin (53) subfamily within the mammalian peroxiredoxin superfamily. Structurally and mechanistically, human PRDX5 exhibits conserved features with bacterial thiol peroxidases (Tpxs) (54). However, multiple sequence alignment guided structural analyses also reveal the same notable divergence: PRDX5 lacks the N-terminal β-flap domain (residues S1-G15) characteristic of bacterial Tpxs. On top of that, PRDX5 has been found to possess an active site proximal extended loop region (amino acid numbers), which folds to form a short helix (Supplementary **Fig. S19**). These unique structural differences between mammalian and bacterial atypical two-cysteine peroxiredoxins may confer unique biochemical features that could be harnessed to design specific structure-guided therapeutic leads. The presence of the unique N terminal β flap region exclusively found in bacterial atypical two-cysteine peroxiredoxin counterparts raises intriguing question about its functional significance, particularly whether it provides adaptive advantages to prokaryotic counterparts in oxidative stress conditions. To address this question, we generated a N-terminal β flap deletion mutant of SaTpx, (NΔ15-SaTpx) lacking amino acids 1–15 and compared its structural and mechanistic roles with the wild type SaTpx through detailed enzyme kinetics, crystallographic and *in vitro* cell based analysis.

### 3.3. Cloning Expression and Purification of WT-SaTpx, NΔ15-SaTpx, SaTrx1 and SaTR

WT-SaTpx, its N-terminal β-flap deletion mutant, R128A-SaTpx and the other proteins of the Staphylococcal thiol-based antioxidant defence pathway, like Staphylococcal thioredoxin 1 (the intermediate electron carrier) and Staphylococcal thioredoxin reductase (SaTR) have been cloned in plasmid-based *E. coli* protein expression vectors (Supp **Fig. S1**). The recombinant proteins were overexpressed under IPTG induction (**Fig. S2**) and for each targeted proteins, purified protein samples were obtained by using immobilized metal affinity chromatography (IMAC using Ni-Sepharose column) followed by size exclusion chromatography. The purified SaTpx and the NΔ15-SaTpx mutant were pre-treated with thrombin protease to remove the N-terminal His tags before the enzymatic assays. Purification of the targeted proteins were monitored through SDS-PAGE analysis and the subunit molecular masses of the target proteins were matched with the SDS-PAGE molecular weight marker ladder (Supp **Fig. S3A, S4A, S5A, S6A**). In-solution oligomeric homogeneity of the purified proteins were ensured by the single sharp 280nm UV absorption peaks obtained after size exclusion chromatography. (Supp **Fig. S3B, S4B, S5B, S6B)**. For biochemical and biophysical assays, the purified proteins were snap frozen by liquid nitrogen and stored at -80^ο^C until further use. For crystallization of WT-SaTpx and NΔ15-SaTpx, freshly prepared protein samples were concentrated to desired concentration ranges using centrifugal membrane filtration technique.

### 3.4. In comparison to WT-SaTpx, NΔ15-SaTpx is catalytically impaired in reducing incoming substrate hydroperoxides

Through the coupled assay it was determined that amongst the three different tested hydroperoxide substrates, tert-butyl hydroperoxide, hydrogen peroxide, and cumene hydroperoxide, tert-butyl hydroperoxide exhibited the highest apparent catalytic efficiency as indicated by its highest apparent K_cat_/K_m_ value (12.8 X 10^3^) compared to two other tested substrates (**Fig. 2c** versus Fig. **S7c and Fig. S7d**). Consequently, the most preferred substrate, tert-butyl hydroperoxide was further used in all the subsequent experiments. Importantly, in presence of the substrate, tert-butyl hydroperoxide, the rate of oxidation of NADPH to NADP+ is significantly reduced in case of NΔ15-SaTpx compared to WT-SaTpx (**Fig. 2a**). Error bars represent standard deviation from triplicate measurements (n=3). The initial velocity of NADPH consumption correlates directly with peroxidase activity, providing a quantitative measure of SaTpx catalytic efficiency in this coupled enzymatic system. The catalytic activity of the purified SaTpx (both WT-SaTpx and NΔ15-SaTpx) was also assessed using Michaelis-Menten (MM) plot by calculating K_m_ and V_max_ values using increasing concentrations of tert-butyl hydroperoxide substrate (**Fig. 2c** and **Fig. 2d**). The obtained enzyme kinetic parameters clearly indicate reduced affinity (as indicated by the K_m_ value) for the substrate in the case of NΔ15-SaTpx compared to WT-SaTpx, whereas the V_max_ value of the reaction remained not significantly reduced. (**Table 1**). The double reciprocal plot has also been plotted to estimate the true kinetic constants and with this analysis we have found that the SaTpx has K_m,_ 20 x10^-6^ M and catalytic efficiency (K_cat_/K_m_) 11x10^4^ M^-1^ s^-1^ while NΔ15-SaTpx has higher K_m_ (59 x10^-6^) and lower catalytic efficiency (1x10^4^ M^-1^ s^-1^) (Figure S9) as compared to wild type enzyme. True kinetic constants have also been reported in Table S3.

**Figure 2:**
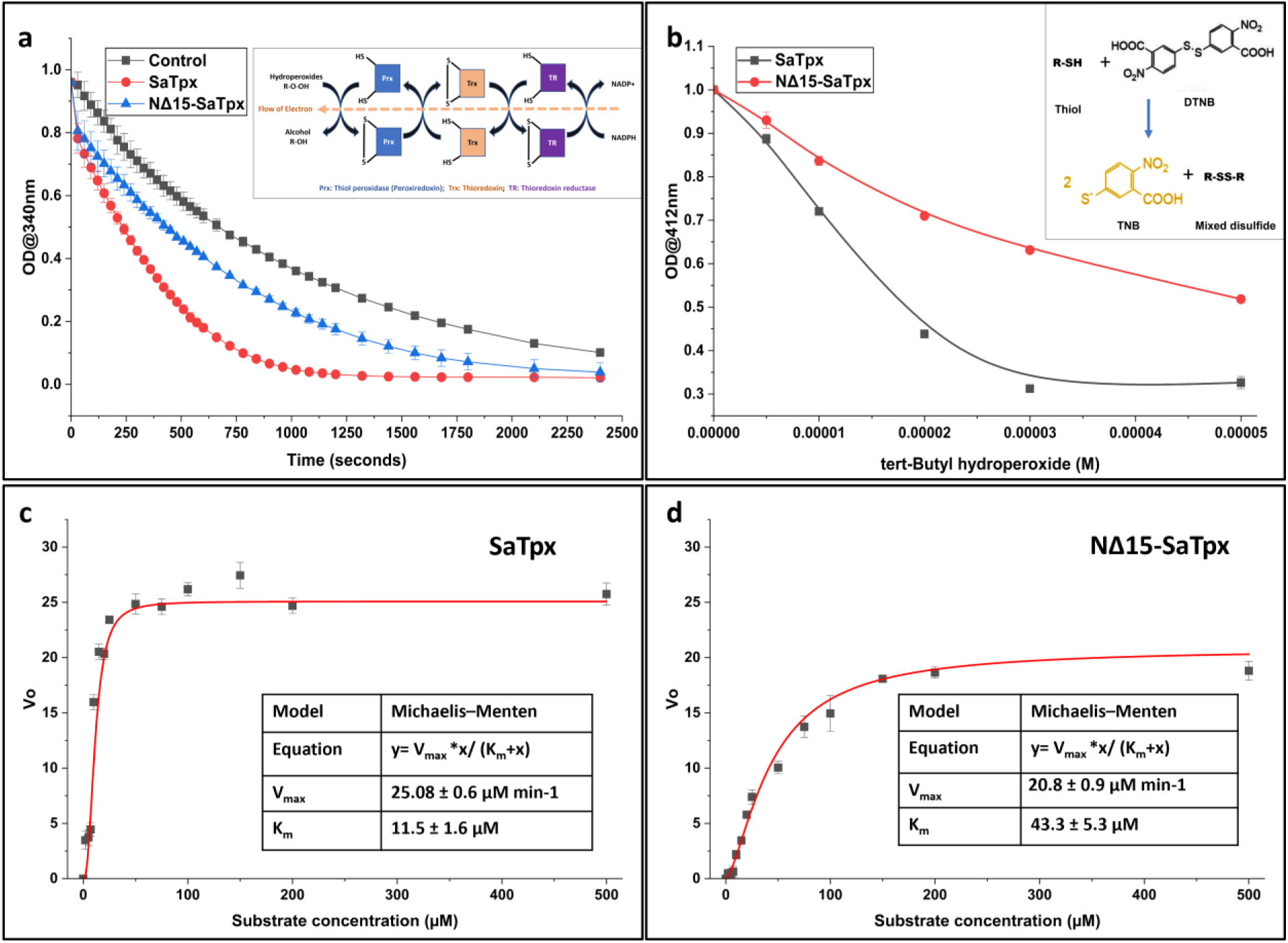
Comparative enzyme catalysis and Kinetic Characterization of Wild-Type and Mutant *Staphylococcus aureus* Peroxiredoxin (SaPrx and NΔ15-SaPrx). **2a**: Time-course analysis of NADPH consumption in a coupled peroxidase assay. Absorbance at 340 nm decreases over time for SaPrx (red), NΔ15-SaPrx (blue), and the control (black-no peroxidase), demonstrating enzyme-mediated NADPH oxidation. **2b: Substrate-dependent oxidation of thiol groups in SaPrx and NΔ15-SaPrx.** The plot shows the decrease in absorbance at 412 nm (OD412) with increasing tert-butyl hydroperoxide (t-BOOH) concentration, indicating oxidation of free thiols. SaPrx exhibits a steeper decline compared to NΔ15-SaPrx, reflecting higher reactivity towards t-BOOH. **2c, d: Michaelis-Menten kinetic analysis of SaTpx (c) and NΔ15-SaTpx (d) with t-BOOH as the substrate.** Initial reaction velocities (V₀) are plotted against substrate concentration and fitted to the Michaelis-Menten equation. SaTpx demonstrates a lower K_m_ (11.5 x10^-6^ ± 0.6M) and higher V_max_ (4.3 x10^-7^ ± 0.06 M sec^-1^) compared to NΔ15-SaTpx (K_m_ = 43.3 x10^-6^ ± 0.9M; V_max_ = 3.5 x10^-7^ ± 0.09 M sec^-1^), indicating higher substrate affinity and catalytic efficiency in the wild-type enzyme. Error bars represent standard deviation from triplicate measurements.

**Table 1:** Comparative apparent enzyme kinetic parameters of SaTpx and NΔ15-SaTpx against tert-butyl hydroperoxide substrates.

| Parameter →<br>Enzymes<br>↓ | K<sub>m</sub><br>(M) | V<sub>max</sub><br>(M sec<sup>-1</sup>) | K<sub>cat</sub> = V<sub>max</sub>/[E]<br>(sec<sup>-1</sup>) | K<sub>cat</sub>/K<sub>m</sub><br>(M<sup>-1</sup> sec<sup>-1</sup>) |
| --- | --- | --- | --- | --- |
| SaTpx | 11.5 x 10<sup>-6</sup> ± 0.6 | 4.7 x 10<sup>-7</sup> ± 0.06 | 9.5 × 10<sup>-2</sup> | 8.3 × 10<sup>3</sup> |
| NΔ15-SaTpx | 43.3 x 10<sup>-6</sup> ± 0.9 | 3.5 x 10<sup>-7</sup> ± 0.09 | 6.9 × 10<sup>-2</sup> | 1.6 × 10<sup>3</sup> |

In this multi-enzyme based coupled reaction, the slower rate of reduction of NADH to NADP+ in the case of NΔ15-SaTpx may indicate its partial inability to reduce the incoming hydroperoxide substrate, which may stem from the reduced affinity of NΔ15-SaTpx towards the incoming hydroperoxide substrate as indicated by the MM plot. However, to precisely ascertain whether NΔ15-SaTpx is partially impaired in catalysing the substrate hydroperoxide reduction, we employed a second reaction strategy, which is single-enzyme based and does not need the functional involvement of SaTrx1 and SaTR proteins.

In the second reaction strategy (i.e. Ellman’s assay), the redox conversion of thiol peroxidase from its free thiol (-SH) containing reduced form to disulfide-linked (-S-S-) oxidized form as a function of increasing concentrations of the substrate t-BOOH has been colorimetrically measured by the addition of DTNB [5, 5’ dithio bis (2-nitrobenzoic acid)]. As the rate of conversion of reduced to oxidized thiol peroxidase increases upon the addition of increasing concentration of the substrate tert-butyl hydroperoxide, the formation of yellow colour of TNB (2-nitro-5 thio benzoic acid; the reaction product of DTNB after interacting with free thiol group) decreases due to the absence of free thiol group of the enzyme. Importantly, the rate of reduction of TNB formation (colorimetrically measured at 412nm) as a function of the substrate tert-butyl hydroperoxide addition may indicate the efficiency of thiol peroxidase to reduce the substrate to the corresponding alcohol (tert-butyl alcohol) and in the process get itself oxidized to DTNB inactive disulfide linked oxidized form. Intriguingly, in contrast to the WT-SaTpx, in the case of NΔ15SaTpx, the rate of reduction of TNB formation upon increasing concentrations of the substrate tert-butyl hydroperoxide addition was found to be significantly reduced (**Fig. 2b**). This result suggests unlike WT-SaTpx, NΔ15-SaTpx has been significantly impaired in reducing the substrate tert-butyl hydroperoxide and, in the process, gets itself oxidized to disulfide-linked form. The inability to get oxidized by the incoming substrate tert-butyl hydroperoxide renders the presence of higher concentration of DTNB active, reduced form of NΔ15-SaTpx, which interacted with DTNB to form more TNB compared to WT-SaTpx.

These results altogether indicate the deletion of the N terminal β-flap region of WT-SaTpx may significantly hamper the catalytic activity of the enzyme.

### 3.5. CD spectroscopy based secondary structure analysis suggests oxidation induced conformational change is preserved in WT-SaTpx and NΔ15-SaTpx

Oxidation induced conformational change at the monomeric level to bring two distantly located cysteine residues (C_P_ and C_R_) to form disulfide bond is common in bacterial thiol peroxidase enzymes (55). In order to assess the three-dimensional structural integrity of NΔ15-SaTpx as well as to follow whether both WT-SaTpx and NΔ15-SaTpx follow the same trivial oxidation-induced conformational changes, we employed CD spectroscopy to determine the secondary structural content of these two enzymes in their reduced and oxidized states, respectively. Importantly, in both of these two enzymes, the CD spectral analysis suggests a characteristic shift of 222 nm **(Fig. S12),** after the addition and incubation with the oxidizing substrate, tert-butyl hydroperoxide. Importantly, in CD spectroscopy of α helix containing proteins, the characteristic peak at 222 nm signifies the relative abundance of the α-helical content. Intriguingly, solution NMR based structural study of *Bacillus subtilis* thiol peroxidase also indicates reduction of α-helical content due to the establishment of disulfide linkage upon the oxidation of the protein. In SaTpx (both in WT-SaTpx and NΔ15-SaTpx) the formation of disulfide bond upon the addition of tert-butyl hydroperoxide substrate has already been indicated in the DTNB based assay **(Fig. 2b)** The CD spectral analysis, on top of it, further suggests the preserved oxidation induced conformational alteration of α helical content of both of these two proteins. Hence, the deletion of the N-terminal β-flap region of WT-SaTpx does not affect the structural integrity of the mutant and the basic mechanism of catalytic turnover mediated through the formation of the disulfide bond.

### 3.6. *In vivo* heterologous production of NΔ15-SaTpx is unable to protect host *E. coli* cells from exogenously induced Reactive Oxygen Species (ROS) stress

Thiol peroxidases are well-known scavengers of exogenously or endogenously produced Reactive Oxygen Species (ROS) (10). Inactivation of thiol peroxidases causes deleterious oxidative inactivation of cellular macromolecules (protein, DNA, membrane lipids etc) which may culminate in the death of the cells(1). *In vitro*, compared to WT-SaTpx, the partial impairment of NΔ15-SaTpx to reduce the incoming substrate hydroperoxides (**Fig. 2** and **Table 1**), led us to test its capability to protect the heterologous host cells from the exogenously induced ROS stress and to compare the same with the WT-SaTpx. *E. coli* host cells harbouring either plasmid borne WT-SaTpx or NΔ15-SaTpx ORFs were exposed to exogenous ROS (tert-butyl hydroperoxide). In parallel, the control groups consisted of *E. coli* host cells containing only the empty plasmid vectors. In each case, IPTG was used to express the cloned enzyme ORFs during ROS induction. In each case, the number of surviving colonies, post-ROS induction, indicates the protective capability of cloned SaTpxs (WT-SaTpx and NΔ15-SaTpx) against exogenously produced ROS. Interestingly, as it has been also observed during *in vitro* enzyme kinetic experiments, heterologous expression of NΔ15-SaTpx is also found to be significantly impaired in its capability to protect *E. coli* host cells, compared to WT-SaTpx. **Fig. 3a** indicates, compared to WT-SaTpx, an approximately three-fold reduction in host *E. coli* cell protection activity of NΔ15-SaTpx against exogenously produced ROS stress at 10 mM tBOOH concentration.

**Figure 3:**
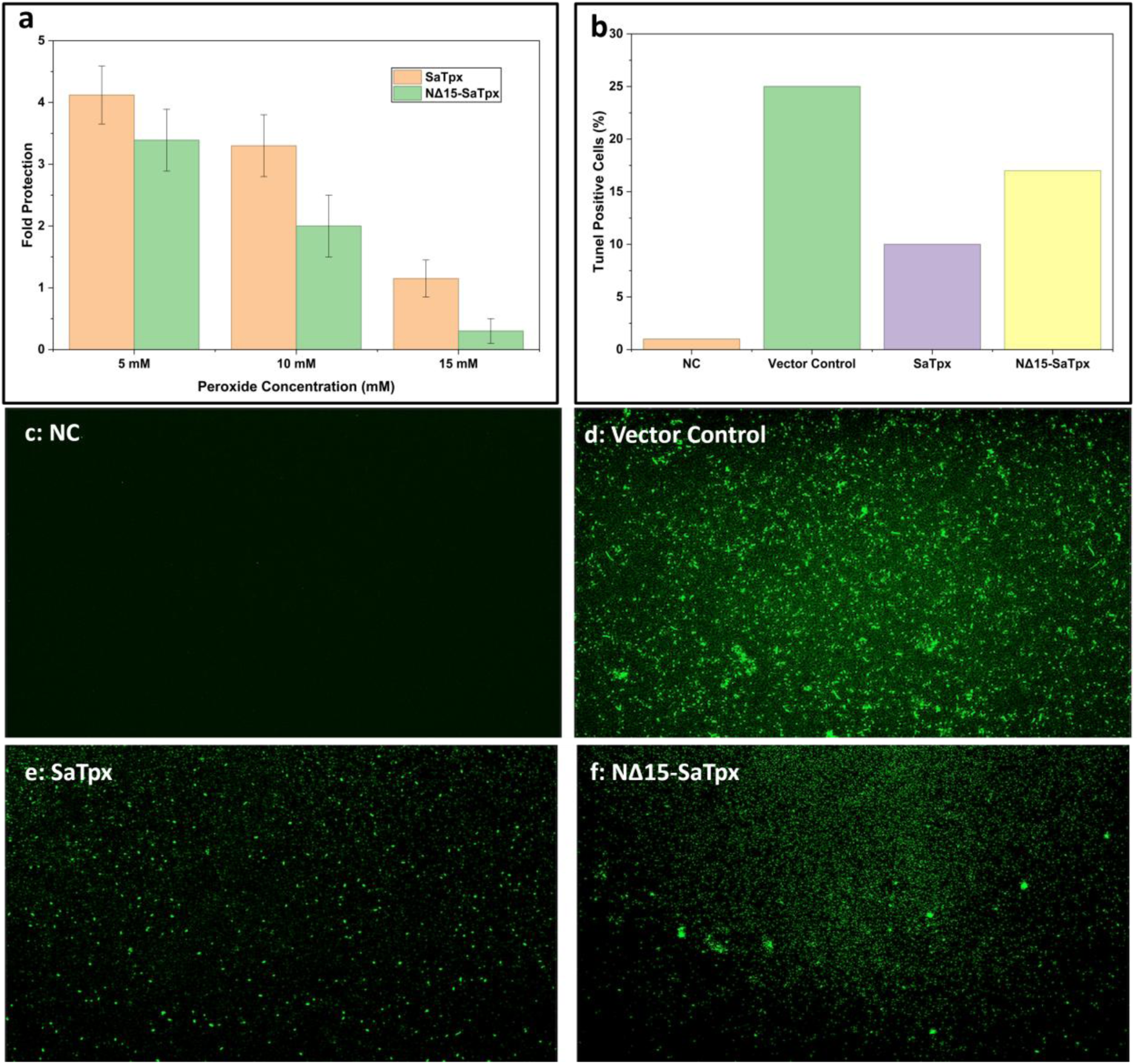
*In vitro* ROS mediated cell and macromolecular (dsDNA) protection activity assessment of SaTpx and NΔ15-SaTpx. **3(a):** *E. coli* cells harbouring recombinant plasmids encoding wild-type peroxiredoxin (SaTpx_yellow) or its N-terminal truncated variant (NΔ15-SaTpx_green) were subjected to tert-butyl hydroperoxide (t-BOOH) stress. Survival was quantified by colony-forming unit (CFU) enumeration at 10⁵-fold dilution. Fold change was calculated by comparing the number of *E. coli* cells with vector control and *E. coli* cells with recombinant gene. Wild-type SaTpx expression conferred significantly enhanced oxidative stress tolerance whereas the protective effect was substantially diminished in the NΔ15-SaTpx variant. Error bars represent standard deviation from triplicate experiments. **3(b)**: Terminal deoxynucleotidyl transferase (TdT) dUTP nick-end labeling (TUNEL) analysis: The bar plot shows the extent of DNA fragmentation. **3(c): Fluorescence micrograph NC** indicates the absence of oxidative stress. **3(d):** Vector control micrograph shows intense fluorescence correlates with maximum DNA fragmentation under oxidative stress**. 3 (e): SaTpx** reduced fluorescence signal compared to vector control indicates that WT-SaTpx limits the DNA fragmentation. **3 (f):** NΔ15-SaTpx has higher fluorescence than WT-SaTpx but less than vector control supports the partial loss of DNA protection in mutant SaTpx.

### 3.7. NΔ15-SaTpx is partially impaired to protect dsDNA break(s) in an *in vitro* metal-catalysed oxidation system

Reactive Oxygen Species and the downstream free radicals generated from their chemical reactions with water molecules are known to break phosphodiester linkages of the nucleic acids (56). Cellular thiol peroxidases are known shield the nucleic acids from the deleterious effects of ROS and their downstream reaction products (57). To assess the *in vitro* phosphodiester bond protection activity of WT-SaTpx and its N-terminal β-flap deleted counterpart (NΔ15 SaTpx), the Metal-Catalysed Oxidation (MCO) protocol was employed to nick supercoiled plasmid DNA in the presence and absence of WT-SaTpx and NΔ15-SaTpx. The MCO generated free radicals attack the phosphodiester linkages of dsDNA and thereby can nick the supercoiled plasmid DNA. The presence of a functional thiol peroxidase (WT-SaTpx or NΔ-SaTpx) was expected to diminish the peroxides generated by MCO system and hence can prevent nicking of the plasmid DNA. The more efficient the enzyme is to scavenge peroxides, the better should be its plasmid DNA nick protection activity. WT-SaTpx was found to protect approximately 55% of the plasmid DNA compared to no-enzyme control reaction. Whereas, in case of NΔ15-SaTpx only 20% of the plasmid DNA was found to be protected from nicking compared to the no-enzyme control reaction (**Fig. S10**). As it is also evident in the enzyme kinetics experiment, the partial impairment of NΔ15-SaTpx to reduce incoming substrate hydroperoxides may also be responsible for its partial impairment to protect plasmid DNA from MCO mediated nicking, which in turn, may also reflect the inability of overexpressed NΔ15-SaTpx to protect host *E. coli* cells from exogenous ROS.

Terminal deoxynucleotidyl transferase (TdT) dUTP nick-end labeling (TUNEL) analysis was executed in order to understand the DNA fragmentation extent in NC (negative control_no treatment), vector control (*E. coli* cells carrying vector pET28a), SaTpx and NΔ15-SaTpx. Figure 3 c, d, e, f represents the micrographs of TUNEL positive cell under various conditions as explained by the histogram (Fig 3b) which represents the quantification of cell death by counting TUNEL-positive cells. *E. coli* cells expressing SaTpx and NΔ15-SaTpx alleviates the H_2_O_2_ effect and provide protection against oxidative stress conditions.

### 3.8. Crystallization and structure solution of SaTpx, NΔ15-SaTpx

So far, compared to the WT-SaTpx enzyme, the NΔ15-SaTpx has been found to show partial impairment in its ability to reduce the incoming hydroperoxide substrates *in vitro* and *in vivo*. The enzyme kinetics data also suggest, NΔ15-SaTpx has reduced affinity for the incoming hydroperoxide substrate compared to its wild type counterpart (WT-SaTpx). In order to decipher the molecular mechanism underlying the observed biochemical differences between WT-SaTpx and NΔ15-SaTpx, we crystallized and solved the crystal structures of these two proteins. The high-quality crystals of WT-SaTpx and NΔ15-SaTpx were obtained using the vapor diffusion hanging drop method. Homogeneously purified WT-SaTpx protein was crystallized with 60mg/ml concentration, with which we got elongated rod-shaped crystals within 24 hours, while for NΔ15-SaTpx, 54mg/ml concentration of purified protein resulted in diamond-shaped crystals after 8 hours of incubation at room temperature. (Supp **Fig. S13**) Both crystal types demonstrated good Xray diffraction capability, yielding well-separated diffraction spots at high resolution (**Table 2**).

### 3.9. Structural features of Staphylococcal atypical 2-Cys peroxiredoxin (SaTpx)

SaTpx is one of the members of the bacterial thiol peroxidase (Tpx) subfamily of enzymes, which also comes under the broader umbrella of two-cysteine peroxiredoxins (Prx). Likewise, the three-dimensional structure of SaTpx is structurally reminiscent of previously reported bacterial Tpxs (RMSD with *E. coli-3hvv 0.941, B. subtilis-2JSZ, 1.093, Streptococcus pneuminoae-1PSQ, 0.628* etc.). Importantly, Prxs are ubiquitous group of proteins present in all three domains of life (**Fig. 1a**). Other than playing the crucial role in organisms’ antioxidative defence pathways, the emerging roles of these proteins as oxidative redox sensors are recently becoming evident. Based on their key cellular role, Prxs are now subdivided as either “sensitive” or “robust” against the incoming organic hydroperoxides (17, 58). Most of the bacterial Tpxs are considered as the “robust” class of Prxs, as their main objective is to mitigate the incoming toxic hydroperoxides, which are often generated by the innate immune cells of the invaded host. Therefore, bacterial Tpxs act as virulence factors, precise inhibition of which attenuates the pathogenic bacteria. Eukaryotic Prxs, however, may also act as the hydroperoxide sensitive counterparts. In the presence of hydroperoxides, reaching a threshold concentration, they become oxidatively inactivated to create an intracellular spatio-temporal concentration gradient of the hydroperoxides as the cellular signalling mechanism (13, 59). Albeit, in terms of overall structure and catalytic mechanisms, bacterial and eukaryotic two-cysteine peroxiredoxins share a high degree of reminiscence, however, prokaryotic Tpxs have a unique N-terminal β-flap region, the function of which is still remained mostly elusive and is highly intriguing (60). In order to delineate the precise structure-function role of the unique N-terminal β-flap region of prokaryotic Tpx, herein, we have determined crystal structures of wild-type Staphylococcal atypical 2-cysteine peroxiredoxin (WT-SaTpx) and the N-terminal β-flap deleted construct of the same enzyme (NΔ15-SaTpx). Furthermore, we also solved the crystal structure of NΔ15-SaTpx construct complexed with a potential tert-butyl hydroperoxide substrate mimic, O-tert-butyl hydroxylamine (OtBH). All of these crystals diffracted to high-resolution limits (1.8Å for WT-SaTpx and NΔ15-SaTpx crystals; 1.55Å for OtBH soaked preformed NΔ15-SaTpx crystals). These high-resolution structural data obtained herein allowed us to unambiguously detect the precise location of the amino acid side chains for the entire range of the SaTpx protomers (Q16-I164) present in the asymmetric unit of the crystal unit cells, the bound substrate mimic (OtBH), the molecules coming from crystallization screen (PEG400) and buffer solutions (DTT). Both WT-SaTpx and NΔ15-SaTpx crystals were indexed, scaled and final structural refinement were carried out in P2_1_2_1_2_1_ space group. However, for OtBH soaked NΔ15-SaTpx crystals the diffraction data were initially indexed in P2_1_2_1_2_1_ space group but was later reindexed, scaled and finally refined in P2_1_ space group due to the presence of pseudo-hemihedral twining as detected by “L-test” (Supp **Fig. S14**). Molecular Replacement phasing method (61) was employed to obtain the initial phases of the diffraction data. The data collection and structure refinement statistics for all the crystal forms have been reported in **Table 2**.

**Table 2:** Data collection and refinement statistics.

| Data Set | WT-SaTpx<br>(PDB Id:<br>8ZWT) | NΔ15-SaTpx<br>(PDB Id:<br>8XVW) | NΔ15-SaTpx-OtBH (PDB Id:<br>8ZXU) | NΔ15-SaTpx-TBH (PDB Id:<br>9XOV) |
| --- | --- | --- | --- | --- |
| Wavelength | 0.97 | 1.54 | 0.97 | 0.97 |
| Resolution range | 33.3 - 1.85<br>(1.916 - 1.85) | 19.14 - 1.803<br>(1.867 - 1.803) | 40.93 - 1.55<br>(1.605 - 1.55) | 36.2 - 1.25<br>(1.295 - 1.25) |
| Space group | P 21 21 21 | P 21 21 21 | P 1 21 1 | P 21 21 21 |
| Unit cell<br>parameters<br>a, b, c (Å) | 50.061,<br>71.911, 88.324 | 38.273, 72.861,<br>148.21 | 38.267, 148.331,<br>72.952 | 38.202 72.392<br>148.145 90 90<br>90 |
| Total reflections | 54414 (4560) | 75172 (6841) | 216264 (21761) | 227484 (22380) |
| Unique reflections | 27447 (2331) | 38875 (3585) | 112479 (11717) | 114567 (11299) |
| Multiplicity | 2.0 (2.0) | 1.9 (1.9) | 1.9 (2.0) | 2.0 (2.0) |
| Completeness (%) | 98.00 (85.44) | 99.08 (93.41) | 99.32 (99.97) | 99.73 (99.86) |
| Mean I/sigma(I) | 13.95 (2.15) | 27.48 (4.31) | 9.03 (3.29) | 12.86 (2.00) |
| Wilson B-factor | 25.95 | 18.57 | 17.5 | 15.59 |
| <sup>a</sup>R<sub>merge</sub> | 0.03084<br>(0.3234) | 0.02215<br>(0.2008) | 0.04786 (0.1471) | 0.02198 (0.327) |
| R<sub>meas</sub> | 0.04362<br>(0.4574) | 0.03132<br>(0.2839) | 0.06768 (0.208) | 0.03109<br>(0.4624) |
| R<sub>pim</sub> | 0.03084<br>(0.3234) | 0.02215<br>(0.2008) | 0.04786 (0.1471) | 0.02198 (0.327) |
| CC (1/2) | 0.999 (0.781) | 1 (0.884) | 0.993 (0.933) | 0.999 (0.819) |
| CC* | 1 (0.936) | 1 (0.969) | 0.998 (0.982) | 1 (0.949) |
| Reflections used in<br>refinement | 27347 (2330) | 38871 (3585) | 116560 (11717) | 114314 (11286) |
| Reflections used<br>for R-free | 1329 (76) | 1945 (174) | 5809 (591) | 5734 (538) |
| <sup>b</sup>R<sub>work</sub> | 0.1697<br>(0.2332) | 0.1564 (0.1945) | 0.1580 (0.2325) | 0.1696 (0.2866) |
| <sup>c</sup>R<sub>free</sub> | 0.2075<br>(0.2851) | 0.1893 (0.2353) | 0.1857 (0.2500) | 0.2004 (0.2809) |
| CC (work) | 0.953 (0.867) | 0.971 (0.910) | 0.917 (0.683) | 0.969 (0.862) |
| CC (free) | 0.957 (0.882) | 0.968 (0.897) | 0.909 (0.625) | 0.950 (0.834) |
| Number of non-<br>hydrogen atoms | 2817 | 2816 | 5479 | 3108 |
| macromolecules | 2543 | 2321 | 4655 | 2486 |
| ligands | 18 | 110 | 343 | 445 |
| solvent | 266 | 385 | 675 | 429 |
| Protein residues | 326 | 298 | 596 | 298 |
| RMS (bonds) | 0.011 | 0.020 | 0.019 | 0.014 |
| RMS (angles) | 1.21 | 1.67 | 2.36 | 1.39 |
| Ramachandran favored (%) | 98.14 | 100.00 | 99.66 | 98.64 |
| Ramachandran allowed (%) | 1.86 | 0.00 | 0.34 | 1.36 |
| Ramachandran outliers (%) | 0.00 | 0.00 | 0.00 | 0.00 |
| Rotamer outliers (%) | 0.00 | 0.78 | 2.15 | 0.72 |
| Clash score | 3.15 | 10.95 | 13.10 | 9.05 |
| Average B-factor | 31.73 | 27.54 | 18.61 | 29.39 |
| macromolecules | 30.80 | 24.40 | 16.93 | 24.51 |
| ligands | 80.85 | 60.16 | 36.16 | 66.47 |
| solvent | 39.15 | 37.10 | 26.34 | 40.97 |
| Number of TLS groups | 1 | 2 | 4 | 1 |

Statistics for the highest-resolution shell are shown in parentheses.

### 3.10. Overall structure

The crystal structure of WT-SaTpx, NΔ15-SaTpx mutant and the substrate mimic (OtBH) bound form of the mutant closely resemble the Prx fold previously reported for other counterparts of the Tpx subfamily. At monomeric level, in each protomer a central layer of twisted β-sheet (β1-β5) is flanked by two layers of α helices (α1, α4 and α2, α3, α5) which is a characteristic feature of the foundational thioredoxin fold with the additional structural elements which are hallmark of Prx fold. Furthermore, the presence of N-terminal β-flap region (Nβ1-Nβ2) in WT-SaTpx structure further justify it is a bona fide member of Tpx subfamily of Prxs. The overall structures of WT-SaTpx, NΔ15-SaTpx and OtBH bound NΔ15-SaTpx were found to be very similar (RMSD values of the pairs: WT-SaTpx/NΔ15-SaTpx=0.24Å, WT-SaTpx/ NΔ15-SaTpx-OTBH= 0.24Å); except for the latter two cases, the deletion of N-terminal β-flap region and the structural changes brought in because of this deletion are noteworthy. At the oligomeric level, both WT-SaTpx and the NΔ15-SaTpx mutant were found to be dimeric, which also substantiates their in solution oligomeric organizations. However, two additional monomers upon a functional dimeric structural organization were found in the asymmetric unit of the pseudo-hemihedrally twinned unit cell of OtBH soaked NΔ15-SaTpx crystals. These two additional monomers also make functional dimeric architecture with the neighbouring monomers of the adjacent unit cells (Supp **Fig. S15**). In all these functional dimeric architectures of SaTpx, the Tpx specific “A type” dimeric interface structure is maintained (17) (**Fig. 4**) which claims approximately 9.7% of the total solvent accessible surface area of the dimer (calculated by PDBePISA (62)). In SaTpx, the formation of this dimeric interface covers total 1423.44 Å^2^ surface area (calculated by Proface (63)). This huge dimeric interface is maintained through the interactions of the amino acid residues principally residing in the loops connecting β2-α3, α4-β4, and β1-α2 as well as the N-terminal amino acid residues of α3 helices of the interacting protomers. Intriguingly, amino acids that stabilize the dimeric interface of Tpx are also found to be highly conserved throughout evolution (64). In SaTpx, out of the total 22 core dimer stabilizing amino acid residues, Threonine 57 (T57) is also one of the most important amino acids of the highly conserved active site catalytic triad of SaTpx. Moreover, the proximity of the other two active site catalytic triad amino acid residues [Cysteine 60 (C60) and Arginine 128 (R128)] (Supp **Fig. S16**) to the SaTpx dimeric interface demonstrates the invincibility of the dimeric interface for the formation of the active site catalytic cleft of SaTpx.

**Figure 4:**
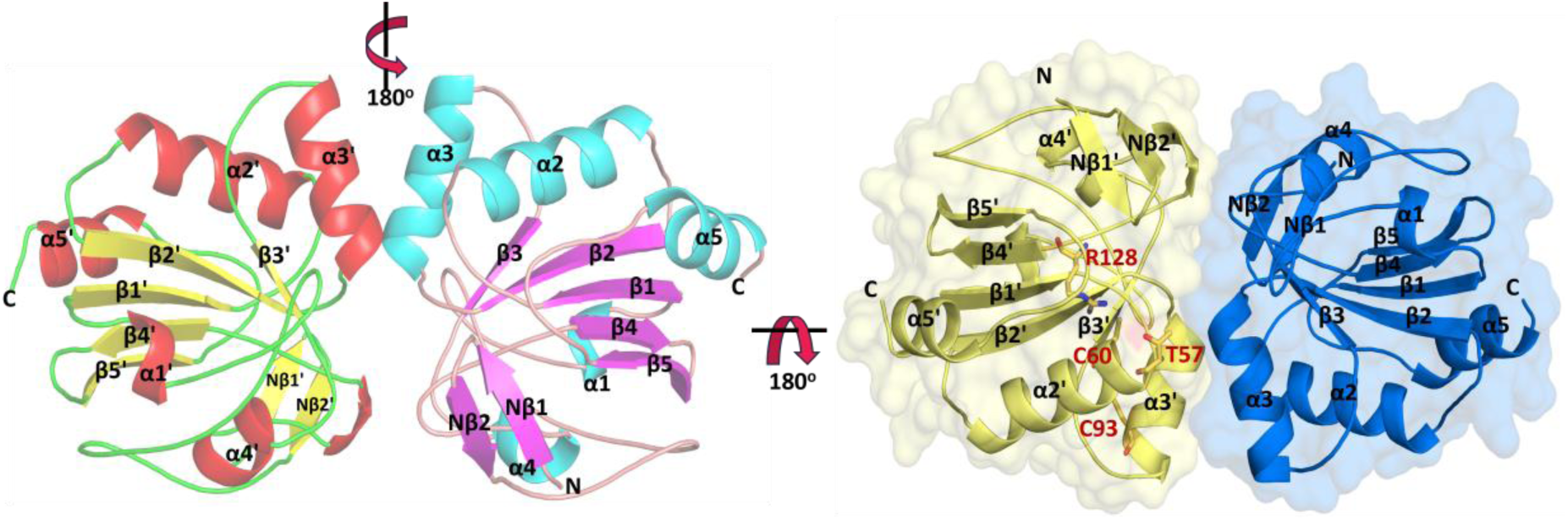
Overall structural feature of SaTpx. Left side cartoon of SaTpx dimer shows the distinct monomers that emphasize the orientation of α-helices and β-strands as well as secondary structural components named. The 180° rotated SaTpx dimer surface and cartoon representations show how the two monomeric units interact mainly through their α3 helices to create the canonical A-type dimer interface forming a stable dimer anchored by hydrogen bonds and hydrophobic interactions. Red highlights the important catalytic residues (Thr57, Cys60, Arg128, and Cys93).

### 3.11. Active sites of WT-SaTpx, NΔ15-SaTpx and OtBH bound NΔ15-SaTpx

#### 3.11.1. WT-SaTpx

The active site of WT-SaTpx resembles a ‘L’ shaped crevice of about ∼36Å length (calculated by MOLE online server (65)), formed at the interface of the two interacting protomers. The active site catalytic triad of SaTpx (comprised of amino acid residues T57, C60 and R128) is situated at the bottom of this crevice. The inner surface of this crevice is mostly guarded by hydrophobic groups of amino acid side chains of two interacting protomers (V52, P53. L125, L123, E122, F6, K7, P10 from one monomer and F87, L34, R108 from the interacting monomer) (**Fig. 5b**), resulting in the high positive logP value (1.2) of this crevice surrounding surface. In typical 2-Cys peroxiredoxins, residues such as Phe, Tyr, Trp, or occasionally Leu/Ile occupy a structural position analogous to the Phe6 found in atypical peroxiredoxins, despite differing sequence locations. A Phe/Tyr around the β1 region occupies a hydrophobic position analogous to atypical Prx positions Phe6. This residue helps stabilize the active conformation, similar to the effect of the β-flap Phe6 in atypical Prx. Unlike the eukaryotic Prxs, this hydrophobic crevice like elongated active site architecture of bacterial Tpxs may favour the binding of lipid hydroperoxides (Supp **Fig. S17**), which also supports the fact that bacterial Tpxs also play the pivotal role of lipid hydroperoxide peroxidase (66). Again, as the name suggests, the catalytic mechanism of two-cysteine peroxiredoxins rely on the presence of two redox active cysteine residues, the peroxidative cysteine (C_P_) and the resolving cysteine (C_R_) (58, 67). In SaTpx crystal structures, these two cysteines (C_P_ and C_R_ are C60 and C93, respectively) are located on α2 and α3 helices, respectively, with a Cα distance of ∼13Å to each other (Supp **Fig. S18**). Absence of any other cysteine residues in the amino acid sequence of the protein unequivocally suggests the sole catalytic redox roles of these two cysteine residues. The absence of formation of a disulfide bond amongst these two cysteine residues and the clear electron density blobs of -Sγ atoms confirm the reduced state of these two cysteine residues which may result from the addition of excess reducing agent (DTT) during the purification of this protein prior to the crystallization trials. Again, the overall structural reminiscence of all crystal forms of SaTpx reported herein (WT-SaTpx, NΔ15-SaTpx and OtBH bound NΔ15-SaTpx) with fully folded (FF) reduced form of *E. coli* Tpx further suggests the fully folded pre-catalytic reduced states of SaTpx crystal structures. In peroxiredoxins, the presence of highly conserved Proline-X-X-X-Threonine/Serine-X-X-Cysteine motif [P(X)_3_T/S(X)_2_C] and one Arginine (R) residue ensures the lowering of pKa value of the peroxidative cysteine (C_P_) present in the motif. Lowering of the pKa of the C_P_ ensue the formation of cysteine thiolate (C-S^-^) anion at physiological pH (7.4), which can further break the peroxide bond of the incoming substrate through nucleophilic attack. The crystal structure of WT-SaTpx shows, C60 may play the role of the C_P_ as its -Sγ atom is in hydrogen bonding proximity with -Oγ atom of T57 (∼3.2Å) and -ηNH2 atoms of R128. (∼3.3Å). Altogether, these three amino acid residues constitute the active site catalytic triad of SaTpx (**Fig. 5a**). In WT-SaTpx active site, while the thiol group (-SH) of peroxidative cysteine, C60 act as a hydrogen bond doner to the -Oγ atom of T57, the resulting partial negative charge on the -Sγ atom of C60 may be stabilized by positively charged -ηNH2 atoms of R128. Moreover, the thiol group (-SH) of C60 is also in hydrogen bonding proximity (∼3.1 Å) of the peptide carbonyl group (>C=O) of S54. This array of hydrogen bonding interactions with the thiol (-SH) group of C60, as well as the close vicinity of the negative charge stabilizing guanidium group [(H2N)(HN)CN(H)-] of R128 renders the lowering of pKa value of C60 thiol group (-SH) and thereby facilitates the formation of the nucleophilic thiolate anion (-S^-^). Importantly, in peroxiredoxins, the highly conserved Arginine (R) and Threonine (T) residues of the catalytic triad are also found to stabilize the incoming hydroperoxide substrate through hydrogen bonding interactions (17, 68, 69). In WT-SaTpx crystal structure, the active site T57 and R128 were found to hydrogen bond two proximal water molecules (W1 and W2 in **Fig 5a**.) which may act like peroxide substrate mimic, Benzoate and acetate at the same position observed in mammalian and mycobacterial counterparts. The evolutionarily conserved Proline residue (P) of P(X)_3_T/S(X)_2_C motif (P53 in SaTpx) provides the structural kink needed to form the active site structural fold as well as provides a hydrophobic shielding of the peroxidative cysteine residue (20, 70, 71). The similar range of low B factors (ADPs) of the catalytic triad residues (ranging from 20-25 Å^2^) suggests a high degree structural stability of WT-SaTpx active site. Importantly, unlike eukaryotic hydroperoxide sensitive Prxs, in bacterial Tpxs the guanidium group [(H2N)(HN)CN(H)-] of the highly conserved Arginine residue has not been observed to adopt multiple conformations upon oxidation and subsequent overoxidation of the sulfhydryl group (-SH) of the peroxidative cysteine (72).

**Figure 5:**
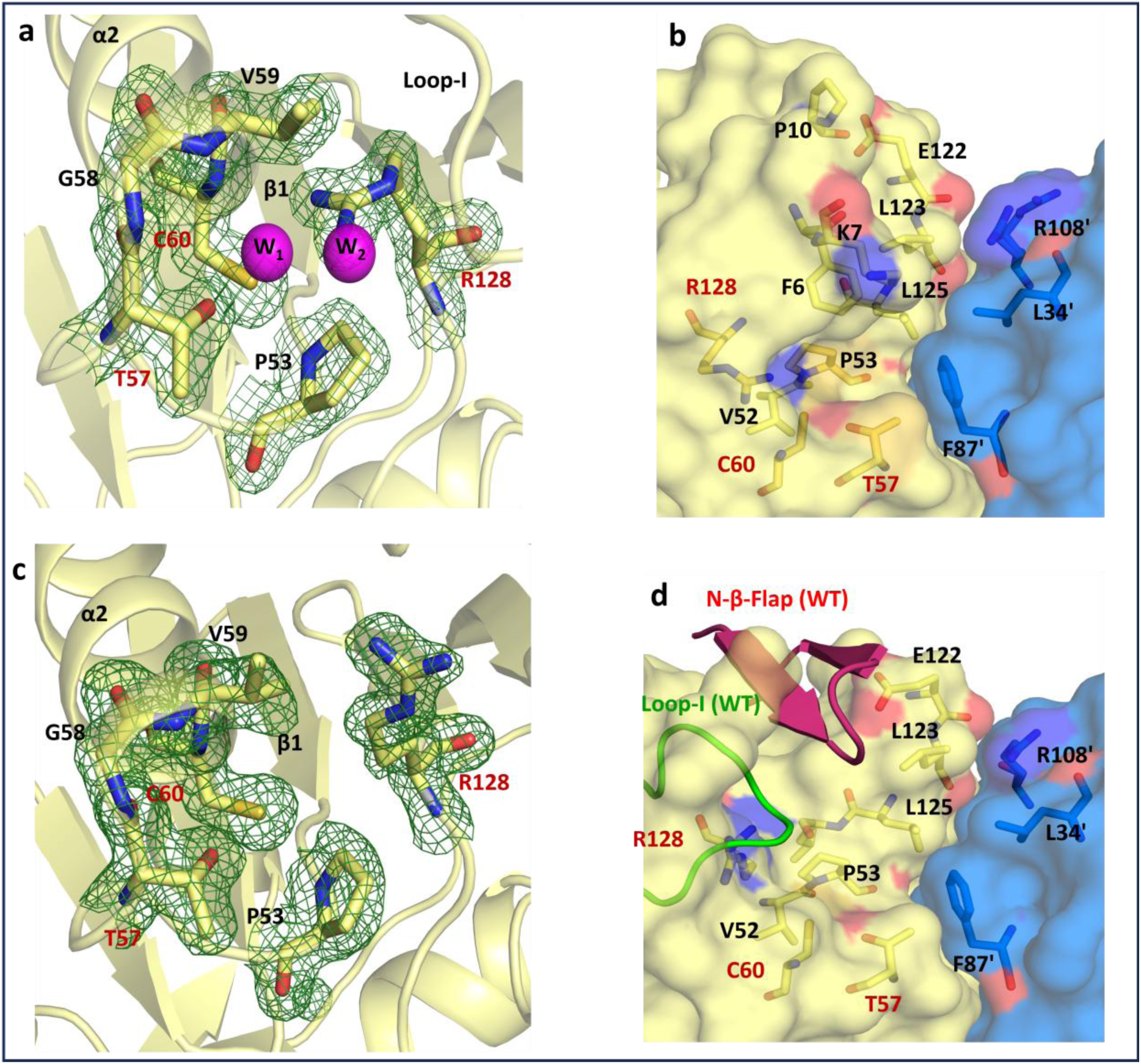
Active site architecture of the targeted proteins (SaTpx and NΔ15-SaTpx). **5a:** Electron density map (2Fo-Fc maps contoured at 1σ) in figure highlight the active site catalytic triad of SaTpx with two water molecules (W1 and W2, magenta spheres) that are positioned inside the catalytic cleft and may resemble the way peroxide substrates bind. The surrounding structural elements (Loop-I, β1, α2) and important catalytic triad residues (Thr57, Cys60, Arg128) are also labelled. **5b:** Surface representation of dimeric interface of SaTpx crystal shows the L-shaped catalytic cleft at the dimer interface. The catalytic triad residues (T57, C60, R128) are positioned at the base of the cleft. Hydrophobic residues such as F6, K7, P10, V52, P53, E122, L123, and L125 from the A chain (yellow) protomer and L34, F87, and R108 from the B chain protomer (blue) primarily line the inner surface of the cleft, forming the substrate-binding environment. **5c:** The SaTpx mutant crystal structure electron density map (2Fo-Fc maps contoured at 1σ) shows the catalytic triad residues T57, C60, R128 (red) as well as Pro53, Gly58, and Val59 residue in proximity of catalytic site within hydrogen-bonding distance. Interestingly, as shown in Fig. 4a, the Arg128 side chain is displaced from its canonical location in the mutant crystal structure of SaTpx. **5d:** Surface representation of the active site of N-terminal deletion mutant SaTpx at its dimeric interface and aligned with wild-type SaTpx (represented in cartoon) highlights the relative position of the N-terminal β-flap (magenta coloured in cartoon representation), which forms the roof of the L-shaped catalytic cavity in the wild-type enzyme. Deletion of the β-flap in the mutant results in a widened cleft. The proximity of Loop-I (green coloured in cartoon representation) to the active site is also evident in wild-type SaTpx, contributing to the compactness of the active site architecture, whereas these features are absent in the mutant, further altering the catalytic environment. Key catalytic and hydrophobic residues at the interface are labelled for reference.

This increased flexibility of the active site Arginine residue, may provide some extra room for the eukaryotic Prxs to accommodate subtle structural changes brought upon by oxidation of the sulfhydryl group (-SH) of peroxidative cysteine, hence, allowing the eukaryotic Prxs to avoid locally unfolded conformation of the active site which acts as a signal for resolving cysteine (C_R_) to attack the oxidized peroxidative sulfhydryl group (C_P_ -SOH) to form the disulfide bond (58). The disulfide-bonded conformation of Prxs is resistant to overoxidation and is the substrate for the downstream electron doner protein thioredoxin. Once again, in eukaryotic Prxs, the hyper flexibility of the active site Arginine residue also provides extra space in the active site. Therefore, in eukaryotic Prxs, after the first round of hydroperoxide mediated oxidation, a second molecule of hydroperoxide can still sit in to ensue overoxidation of the preformed sulfenic acid (C_P_-SOH) of the peroxidative cysteine to form cysteine sulfinic acid (C_P_-SO_2_H). Hence, increased flexibility of the guanidium group of the active site Arginine residue of eukaryotic Prxs may account for their augmented propensity for overoxidation, thus making them less robust compared to the bacterial Tpxs. In contrast to that, the guanidium group of active site Arginine residue of bacterial Tpxs is firmly held at its catalytic position by tight steric packing of active site proximal loop-I region (V145 to F151 in WT-SaTpx) which is further stabilized by extensive arrays of hydrogen bonding interactions with the unique but highly conserved N-terminal β-flap region of prokaryotic Tpx (M1 to L13 in WT-SaTpx) (**Fig. 6a**.). The lesser flexibility of the active site Arginine side chain in bacterial Tpxs may not support overoxidation of C_P_SOH, by not allowing a second molecule of hydroperoxide to sit in the Tpx active site, as well as by swiftly transducing the peroxidative oxidation induced conformational signalling to allure the resolving cysteine (C_R_) to attack and form intramolecular disulfide bond. In order to assess the plausible active site Arginine stabilizing role of the highly conserved N-terminal β-flap region of bacterial Tpxs, initially through a molecular dynamics simulation experiment, we tested the altered flexibility of R128 residing loop of SaTpx as a function of the presence or absence of amino acids forming the N-terminal β-flap region of WT-SaTpx (M1-G15). Intriguingly, as indicated by post-MD simulation, Root Mean Square Fluctuation (RMSF) analysis of Cα trails of these two constructs [WT-SaTpx versus NΔ15-SaTpx (where we manually deleted β-flap region)] we found, indeed, the selective deletion of the N-terminal β-flap region of SaTpx, increases the flexibility of R128 bearing loop of SaTpx (E121 to A129) with a significant structural change through loop-I region (V145 to F151 in SaTpx) of the protein (**Fig. 6b**). This observation further instigated us to set out for the structure-guided functional characterization of N-terminal β-flap deletion mutant of SaTpx, to uncover the yet unexplored role N-terminal β-flap region of prokaryotic Tpxs.

**Figure 6a:**
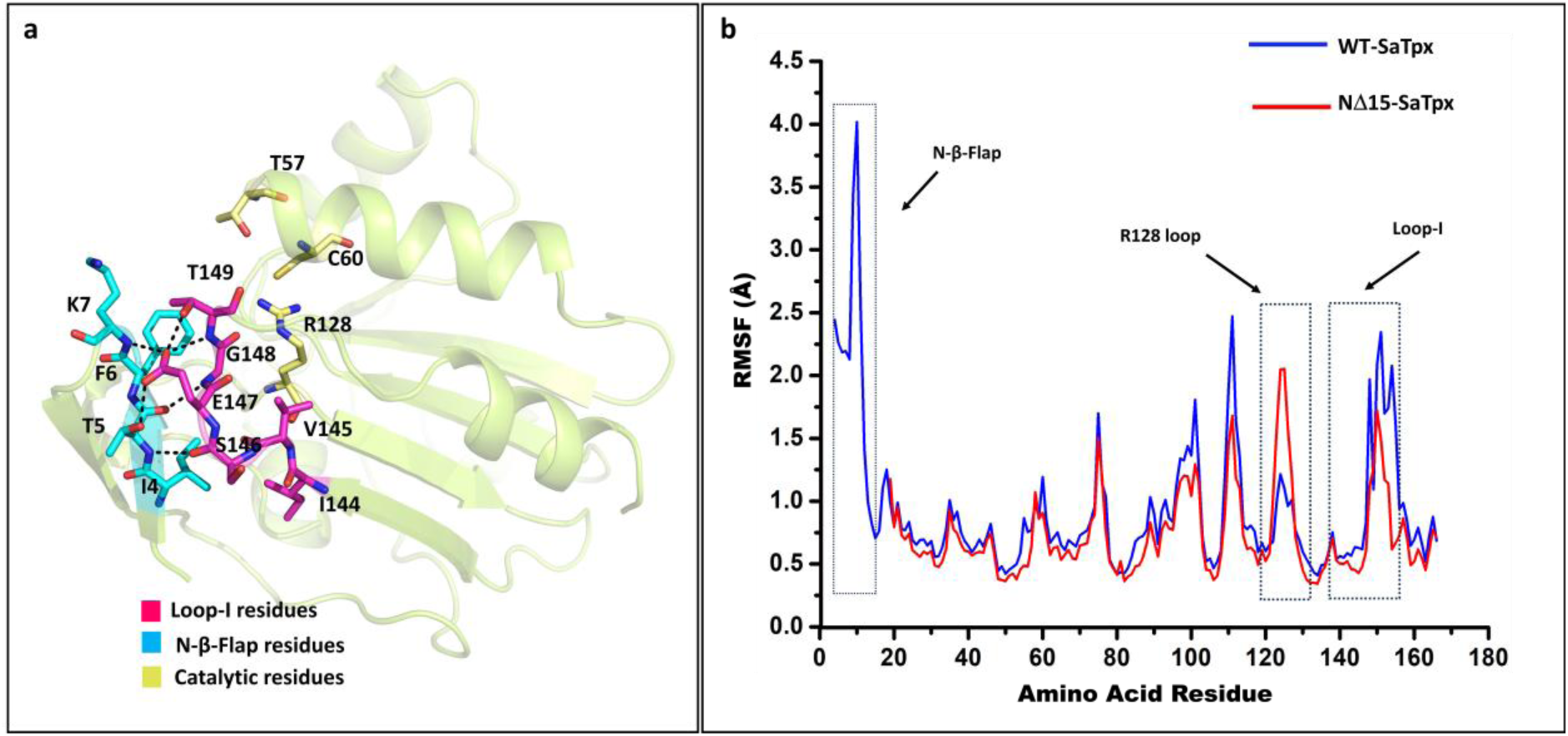
N terminal β sheet stabilizes the active site proximal loop-I region which is further needed to stabilize the catalytic orientation of Arg128. Cartoon representation of wild-type SaTpx highlighting the spatial arrangement of Loop-I residues (magenta-sticks), N-terminal β-flap residues(blue-sticks), and catalytic residues (yellow-sticks). Stick models illustrate extensive hydrogen bonding between N-β-flap residues and both Loop-I residues and catalytic amino acids, including Arg128. This network of interactions stabilizes the active site architecture. **6b: MD simulation Based assessment of the role of N terminal β sheet of SaTpx in regulation of its catalytic activity:** The residual fluctuation of amino acids over the MD simulation time trajectory has been analysed by comparing the RMSF profiles of SaTpx (represented by blue line) and the NΔ15-SaTpx (represented by red line) According to the plot, both proteins exhibit greater flexibility in the Arg128 containing loop region, while the mutant exhibits somewhat higher fluctuations in this region. On the other hand, the mutant Loop-I region shows lower RMSF values than the wild type SaTpx which corroborates the crystal structure data and suggest that the deletion of the N-terminal β-flap has affected the Loop-I dynamic behaviour and Arg128 conformational flexibility.

#### 3.11.2. NΔ15-SaTpx

Importantly, other than the absence of the N-terminal β-flap region and the structural alteration associated with that deletion, the overall structure of NΔ15-SaTpx remains mostly unaltered at the dimeric and monomeric levels compared to the WT-SaTpx. Amongst these two counterparts, the monomeric RMSD value of 0.24 Å and dimeric RMSD value of 0.32Å exemplify this fact. However, enzyme activity data shows, in contrast to the WT-SaTpx, NΔ15-SaTpx is significantly catalytically impaired in part. The comparison of the active site catalytic cleft of these two variants shows, unlike WT-SaTpx, the catalytic cleft of NΔ15-SaTpx becomes shallow and much widened (**Fig. 5d**). Intriguingly, structural superimposition of the active sites of these two variants suggests, this widening not only caused by the absence of the N-terminal β-flap region (which has been found to form the collar of WT-SaTpx active site crevice) but also resulted from the displacement of the active site proximal loop-I region from its original position by ∼ 6.5Å (based on relative Cα positions of T149 in these two constructs) (**Fig. 5c**). Importantly, as it could be also predicted through the MD simulation results, concomitant with the structural displacement of loop-I region, the side chain guanidium group of the R128 residue was also found to be displaced by ∼ 4Å from the catalytic centre of NΔ15-SaTpx, resulting in flipping away of -ηNH1 and -ηNH2 atoms of R128 by ∼37^ο^ and 42^ο^ respectively from the NΔ15-SaTpx catalytic centre. However, unlike R128, the positions of other two highly conserved amino acid residues of SaTpx catalytic triad (C60 and T57) were found to be unaltered in NΔ15-SaTpx. In NΔ15-SaTpx, the flipping of R128 guanidium group disengages itself from the hydrogen bonding interaction with -Sγ atom of the peroxidative cysteine C60 (**Fig. 7b**). The loss of R128 guanidium group mediated charge coupled stabilization interaction of peroxidative cysteine thiolate (Cys-S^-^) may account for the significantly reduced catalytic efficiency of NΔ15-SaTpx. Therefore, in NΔ15-SaTpx mutant, the loss of the Tpx specific N-terminal β-flap region caused the differential structural disposition of loop-I region, which resulted in the increased flexibility and flipping of the R128 residue **(Fig. 7a**) of the active site catalytic triad, further culminating in the reduced catalytic efficiency of the mutant protein.

**Figure 7:**
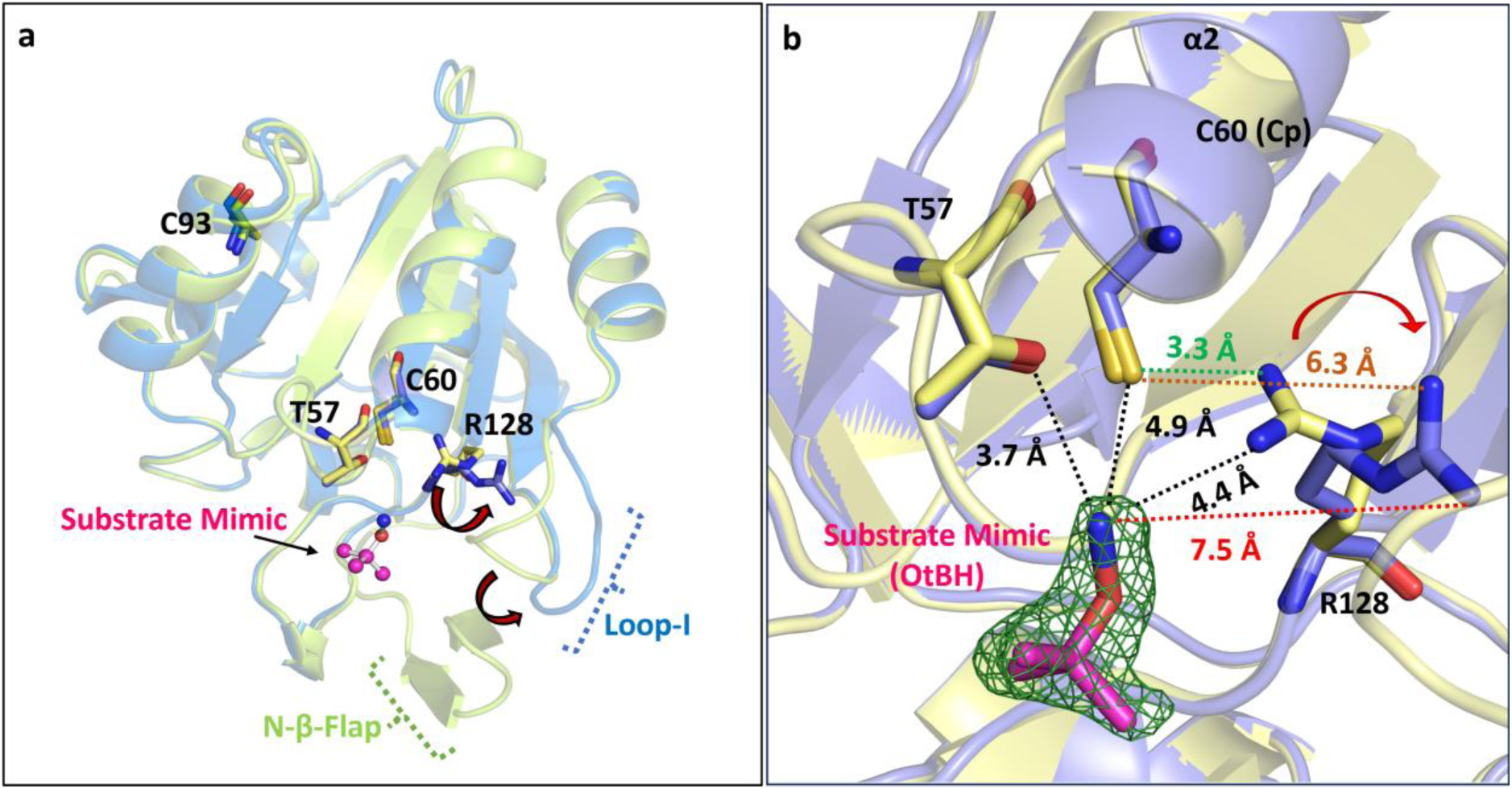
Structural alignment of wild-type *Staphylococcus aureus* peroxiredoxin (WT-SaTpx, green) with the OTBH-bound NΔ15-SaTpx mutant (blue). Figure 7a illustrates the spatial arrangement of key active site residues (T57, C60, R128) and the substrate mimic OtBH (magenta). **7b**: The right panel highlights the displacement of the Arg128 guanidinium group: in NΔ15-SaTpx, the ηNH₂ atoms of Arg128 are positioned 7.5 Å from the NO6 atom of OtBH, compared to 4.4 Å in WT-SaTpx, indicating a loss of stabilizing hydrogen-bonding interactions with the substrate analog due to β-flap deletion. Distances between substrate mimic, catalytic residues, and key conformational shifts are annotated.

Interestingly, R128A point mutant of SaTpx shows complete loss of enzymatic activity (Supplementary S8a, b Figure). In comparison to that NΔ15-SaTpx shows partial loss of hydroperoxidase activity. In this context, in order to solve this ambiguity, we have solved an ultra-high resolution crystal structure of NΔ15-SaTpx (data collection and refinement statistics reported in Table2) which allows us to detect high-resolution structural information and multiple alternative conformations of amino acid side chains present in the NΔ15-SaTpx protein. Intriguingly, we found in this structure, the guanidium group [(H2N)(HN)CN(H)-] containing side chain of active site R128 adopts a dual conformation. In one conformation guanidium group [(H2N)(HN)CN(H)-] of R128 is close to the thiol group (-SH) of C60 and making hydrogen bond (3.1 Å), in another conformation, the side chain of R128 flips away resulting in the loss of the hydrogen bonding with the thiol group of C60 (Fig S20). Both of these two alternative conformations of R128 has been refined to 50% occupancy. Therefore, the deletion of the highly conserved N-terminal beta flap region of SaTpx confers increased structural flexibility of R128 side chain which may cause the reduced catalytic efficiency of this mutant; rather than the complete loss of the catalytic activity as it has been observed in Arg128A point mutant.

Compared to WT-SaTpx, the reduced catalytic efficiency of NΔ15-SaTpx has been also found to be corroborated with its significantly reduced ability to protect purified plasmid DNA from peroxides generated through Fenton reaction *in vitro* **(Fig. S10**); as well as its inability to protect the *E. coli* expression host from the tert-butyl hydroperoxide induced oxidative stress under IPTG induction (**Fig. 3a**).

Importantly, mammalian peroxiredoxin 5 (PRDX5) is the unique member and the solo representative of atypical two-cysteine peroxiredoxin sub-family amongst the six other reported family members of eukaryotic peroxiredoxins. Structurally and functionally PRDX5 is also highly reminiscent with bacterial Tpxs and just like bacterial Tpxs is more resistant against active site thiol hyper oxidation. Surprisingly, PRDX5 also lacks the N-terminal β-flap region of bacterial Tpxs; nonetheless, unlike the NΔ15-SaTpx, the loop-I region of PRDX5 is still maintained at the close proximity of the active site catalytic triad (Supp **Fig. S19**). Intriguingly in PRDX5, the loop adjoining α4 and β6 got extended and folded into a short helix, α5 (Supp **Fig. S19**). This extended helix is unique for PRDX5 and participates in the formation of the active site cleft of the protein (53) and may also play the analogous role of bacterial Tpx specific N-terminal β-flap region to stabilize the loop-I region and in turn, resist the swinging of active site Arginine residue from its catalytic position. Interestingly, the flipping of the active site Arginine residue has been also previously reported in couple of other peroxiredoxin subfamilies like Peroxiredoxin Q from *X. campestris* (PDB Id: 5INY (73)), and in human Prx6 (PDB Id: 1PRX (74)); but in both of the cases, the swinging of the active site Arginine resulted from the hyperoxidation of the peroxidative cysteine (C_P_). In these cases, even after soaking with the substrate hydroperoxides, due to crystal lattice constrains these proteins could not undergo peroxidative cysteine oxidation mediated partial local unfolding (PLU) of the active site which would otherwise have generated the signal to form disulfide bonded oxidized conformations(75). However, both of these Prxs, do not belong to the Tpx family of ‘robust’ peroxiredoxins and lack the N-terminal β-flap region. Moreover, soaking the preformed SaTpx (both WT and NΔ15) crystals with its substrate organic hydroperoxides, always resulted in melting of the crystals with complete loss of the X-ray diffraction capability. In SaTpx, the two redox active cysteines [C_P_(C60) and C_R_(C93)] are ∼13Å apart from each other in their fully reduced states. A huge conformational change with dissolution of few secondary structural elements of the reduced state conformation is needed to bring these two cysteines close together to form disulfide bonded oxidized state of the protein (Supp **Fig. S12d**). Soaking of the preformed SaTpx crystals (even the partially active NΔ15-SaTpx mutant crystals) with the prevailing reduced conformation of the protein with substrate hydroperoxides, hence may disrupt the existing lattice contacts resulting in complete loss of the X-ray diffraction. Again, other than facilitating the formation of peroxidative cysteine thiolate anion (C_P_-S^-^), prior to catalysis, the conserved Arginine residue at the catalytic triad of Prxs also have been found to stabilize the incoming substrate by formation of specific hydrogen bonds (14). Hence, flipping of the side chain guanidium group of the R128 of NΔ15-SaTpx may also hinder its substrate binding capabilities. Compared to WT-SaTpx, in NΔ15-SaTpx the reduced affinity (Table 1) (as determined by the K_m_ value) for the substrate, tert-butyl hydroperoxide may also reflect this fact.

#### 3.11.3. OtBH bound NΔ15-SaTpx

Till date, the substrate bound precatalytic state conformation of Tpxs is not available. Moreover, in order to delineate the impact of structural disposition of R128 guanidium group on substrate binding, we soaked the preformed WT-SaTpx, and NΔ15-SaTpx crystals with the substrate, tert-butyl hydroperoxide as well as with structural mimic of tert-butyl hydroperoxide [O-(tert-butyl) hydroxylamine (OtBH)]. Although none of the crystals of WT-SaTpx, and NΔ15-SaTpx showed any diffraction after soaking with tert-butyl hydroperoxide, brief (15min) soaking of NΔ15-SaTpx crystals with the substrate mimic OtBH resulted in high quality diffraction, yielding high resolution difference density maps of the OtBH molecules bound at the active site of NΔ15-SaTpx (**Fig. 8**). Unlike the active site of WT-SaTpx where two water molecules were found to occupy the substrate binding site and may mimic the substrate hydrogen peroxide (H-O-O-H), the active site of NΔ15-SaTpx is occupied with PEG400 molecules which came from the crystallization buffer. The brief soaking of O-(tert-butyl) hydroxylamine (OtBH) to the preexisting crystals of NΔ15-SaTpx resulted in displacement of PEG400 molecules from the active site of NΔ15-SaTpx protomers. Amongst the four protomer chains found in the asymmetric unit of hemihedrally twinned OtBH soaked NΔ15-SaTpx crystals, the active sites of chain A and B were still found to be occupied with PEG400 **(Fig. 8c(I), Fig. 8c(II))** molecules while chain C represents exclusive presence of a single OtBH molecule (**Fig. 8c(III))** and chain D demonstrates the presence of two OtBH molecules along with a preexisting PEG400 molecule **(Fig. 8c(IV)).** The occupancy of the bound OtBH ligands were refined to 100%, justifies their stable binding at the respective binding sites within asymmetric unit of the crystal lattice. The active site bound PEG400 molecules may represent a weak substrate mimic as it has been found to project its terminal hydroxyl groups [HO-(CH_2_-CH_2_O)_n_-H] towards the active site catalytic cleft of NΔ15-SaTpx. The selective replacement of PEG400 molecules by OtBH and its exclusive positioning at the NΔ15-SaTpx active site may justify its structural mimicry with the substrate, tert-butyl hydroperoxide. Once again, soaking of WT-SaTpx crystals with OtBH resulted in rapid crystal melting with no diffraction. The superimposition of WT-SaTpx active site structure with the active site of NΔ15-SaTpx, where one OtBH molecule is exclusively positioned (chain C), divulges, for the first time, the important structural insights of substrate binding to the active site catalytic cleft of Tpxs. Furthermore, it also clarifies the indispensable incoming substrate stabilizing role of the highly conserved, active site catalytic triad of Tpxs. Again, the novel structural role of Tpx specific N terminal β-flap region to regulate the catalytically important structural disposition of the active site Arginine residue could be also envisaged from this structural comparison. Superimposition of the active site of WT-SaTpx with the OtBH bound active site of NΔ15-SaTpx suggests the plausible substrate binding sites. The OtBH soaked crystal structure of NΔ15-SaTpx defines two separate binding sites of the small organic hydroperoxides to reach the peroxidative cysteine residue (C60). In different protomers of OtBH soaked NΔ15-SaTpx crystal both of these sites have been found to be contested between the preoccupied PEG400 molecule and the soaked incoming OtBH molecules. However, amongst these two sites, the one (observed in the C chain of OtBH bound NΔ15-SaTpx crystal), which allows the projection of the NO6 atom of the incoming OtBH molecule to the -Sγ atom of peroxidative cysteine (C_P_) by placing the OtBH molecule in close proximity to possible hydrogen bonding interactions with T57 (3.7Å) and R128 (4.4Å) residues in WT-SaTpx active site, seemed to be the most plausibly optimal precatalytic positioning of the bound substate molecule. In this state, the tert-butyl group of the bound OtBH molecule faces the hydrophobic inner wall (guarded by hydrophobic sidechains of F6, K7, P53, M120, L123, L125 T149 of one protomer and L34, F87 of the interacting protomer) of the active site catalytic crevice of WT-SaTpx while its O-N bond mimicking the O-O bond of the substate hydroperoxide positions itself at the bottom of the L-shaped active site where the conserved amino acids of the active site catalytic triad (T57, C60, R128) reside. Compared, to the superimposed WT-SaTpx active site, in NΔ15-SaTpx, the swinging of R128 side chain resulted in the increased distance of -ηNH2 atoms of R128 and the NO6 atom of the incoming OtBH molecule (from 4.4Å to 7.5Å) **(Fig. 8c(III)).** This may culminate into loss of a possible hydrogen bonding interaction to stabilize the incoming substrate in the NΔ15-SaTpx active site pocket. This position of bound OtBH in NΔ15-SaTpx crystal, however would supposed to be a bit restrained in WT-SaTpx, as the side chain phenyl ring of the active site guarding F6 residue is making a short contact with the CO4 atom of the tert-butyl group of bound OtBH molecule bound in NΔ15-SaTpx crystal (which lacks the F6 bearing N terminal β-flap region). Intriguingly compared to the WT-SaTpx, this slight off-positioning of the substrate mimicking OtBH molecule in NΔ15-SaTpx active site may stem from the loss of hydrogen bonding interaction with R128 guanidium group due to its flipping from the catalytic centre of NΔ15-SaTpx. This possible loss of incoming hydroperoxide substrate stabilizing hydrogen bonding interaction in NΔ15-SaTpx may result in the reduced affinity for incoming substrates, which has also been reflected by the increased K_m_ of NΔ15-SaTpx compared to WT-SaTpx, against tert-butyl hydroperoxide substrate (**Table 1**).

**Figure 8:**
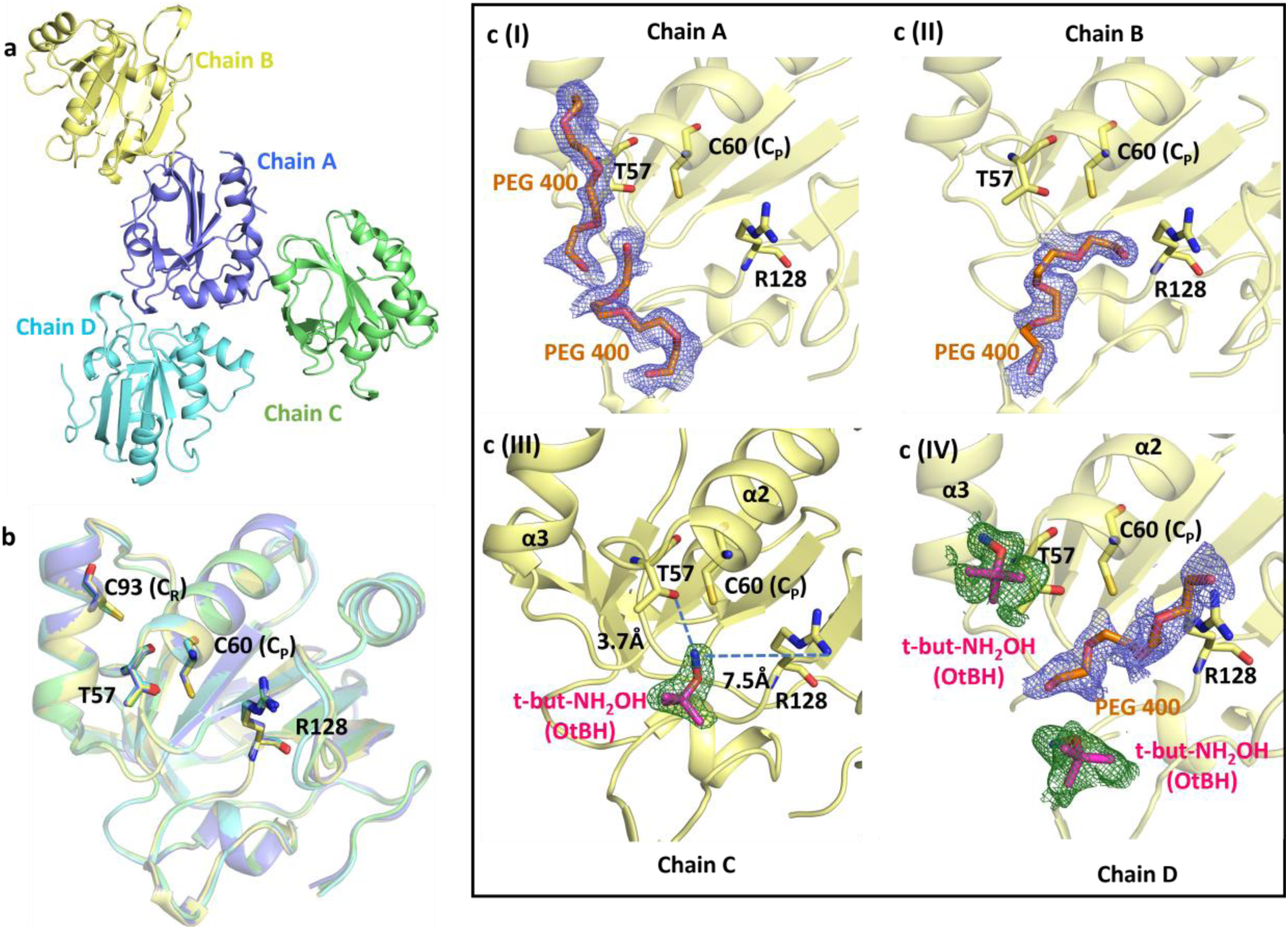
NΔ15-SaTpx-OtBH bound crystal structure protomers. **8a:** Crystal structure of OtBH-bound NΔ15-SaTpx reveals four protomers (chains A–D) in the asymmetric unit. **8b:** Structural alignment of the NΔ15-SaTpx and OtBH-bound NΔ15-SaTpx demonstrates no significant conformational changes upon substrate mimic binding. **8c: Electron density maps highlight ligand binding in each protomer: (c[I])** chain A accommodates two PEG400 molecules near the active site, **(c[II])** chain B binds one PEG400 molecule at the catalytic cleft, **(c[III])** chain C features OtBH positioned with its NO6 atom oriented toward the Sγ atom of the peroxidatic cysteine (C60), enabling potential hydrogen bonding with T57 (3.7Å) and R128 (4.4Å), **(c[IV])** chain D contains both PEG400 at the active site and OtBH adjacent to it. The observation of PEG400 and OtBH binding in different protomers supports the existence of dual substrate-binding loci in proximity to the active site of NΔ15-SaTpx.

### 3.12. Plausible Role of N terminal β-flap region in Tpx subfamily of Prxs

The presence of the N terminal β-flap region is one of the hallmark structural features that all members of Tpxs possess. Once again, Tpxs represent the major subfamily of ubiquitously obtained atypical two-cysteine peroxiredoxin family of proteins. Interestingly, irrespective of high degree of structural and functional reminiscence, eukaryotic atypical two-cysteine peroxiredoxins lack the N terminal β-flap region like bacterial Tpxs. Despite of decades of research efforts to decipher the structure guided functional attributes, and catalytic mechanism of Tpxs, the precise role of the N terminal β-flap region is obscured as yet. Tpxs are robust mitigators of host-induced peroxide stress and play major role in bacterial pathogenesis. The structural differences of Tpxs with their eukaryotic counterparts can be harnessed to design pathogen-specific inhibitor leads which can selectively inhibit Tpxs. In the present work, we determined the high-resolution crystal structures of WT-SaTpx (Tpx ortholog from *Staphylococcus aureus*), as well as NΔ15-SaTpx (Staphylococcal Tpx ortholog lacking the N terminal β-flap region) and OtBH bound NΔ15-SaTpx (NΔ15-SaTpx bound to the substrate analog O-tert-butylhydroxylamine) to pinpoint the role of the N terminal β-flap region of Tpxs. The high-resolution crystal structures of the aforementioned protein constructs provided us the detailed structural glimpse of the highly orchestrated catalytic triad of WT-SaTpx (constituted by T57, C60 and R128 residues) as well as also shows the disruption of the same catalytic triad in case of NΔ15-SaTpx. The disruption of the catalytic triad is caused by swinging of - R128 side chain guanidium group [(H2N)(HN)CN(H)-] away from the thiol group (-SH) of peroxidative cysteine (C60), resulting in the loss of hydrogen bonding interaction between ηNH2 atoms of R128 and -Sγ atom of C60. The catalytic mechanism of thiol peroxidases essentially depends on the formation and stability of the nucleophile, thiolate anion (-S^-^) of the peroxidative cysteine, which eventually attacks and cleaves the peroxide bond (O-O) through SN2 mechanism. The lowering of pKa value of the C60 thiol group is ascribed by the hydrogen bonding vicinity of T57 and R128. Oγ1 atom of T57 draws the proton from thiol group of C60, thereby forming the cysteine thiolate anion (-S^-^) nucleophile, which is also stabilized by nearby positively charged guanidium group of R128. On the other hand, the same positively charged guanidium group of R128 can also act as the Lewis acid to stabilize the peroxide bond of the incoming substrate, increasing the electrophilicity of the bound substrate for the subsequent attack by the C60 thiolate nucleophile. (**Scheme 1**). Hence in the active site of NΔ15-SaTpx, the swinging of the guanidium group of R128 has multi-fold effect to render the mutant enzyme catalytically inefficient. The altered conformation R128 side chain in NΔ15-SaTpx was refined to 100% occupancy and the same range of average B factor of the side chain atoms of R128 with the neighbouring atoms also fortifies its stable conformational disposition.

**Scheme 1:**
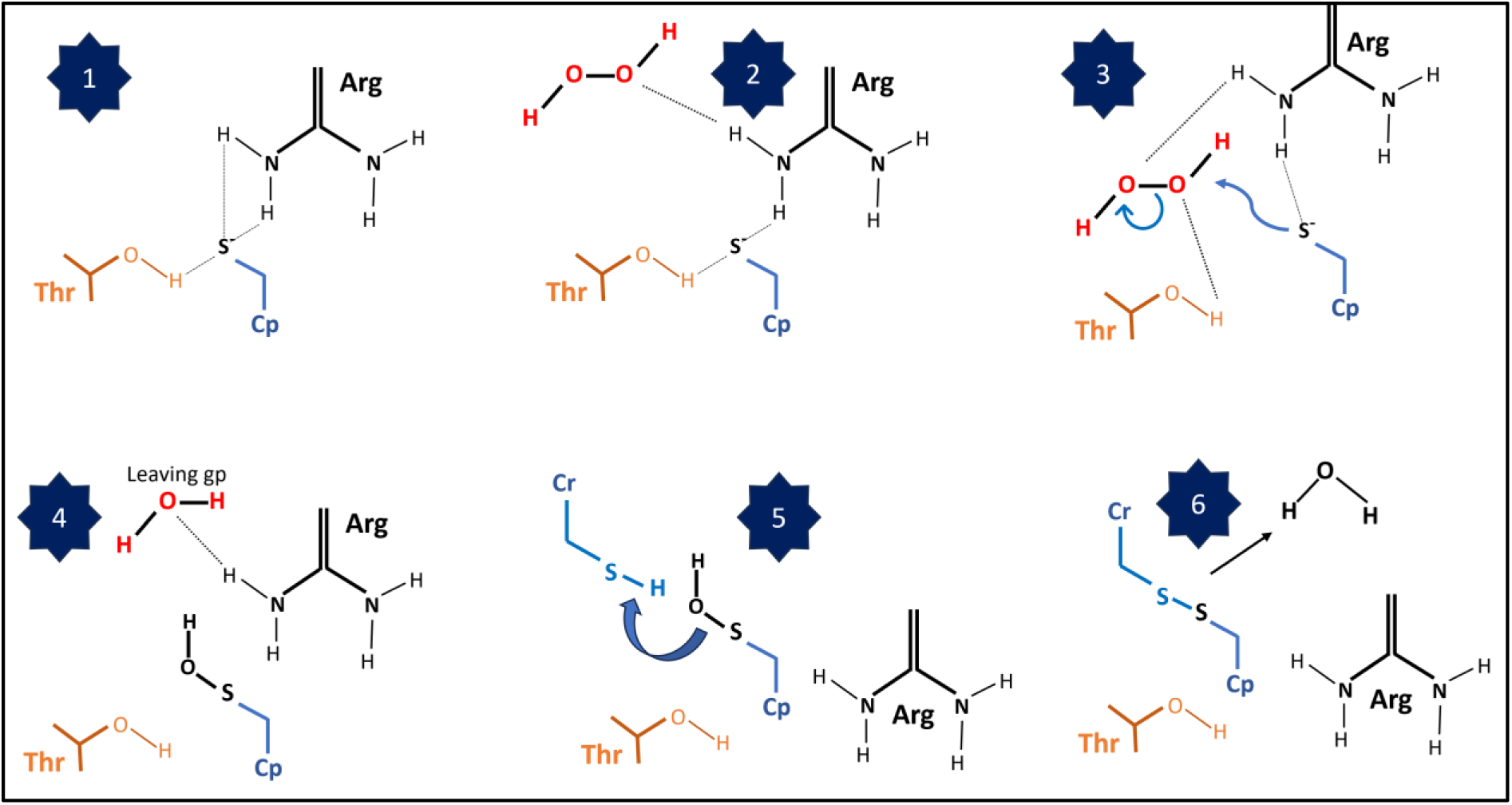
Molecular interplay and substrate dissociation reaction of SaTpx: In the given scheme the **1.** panel shows that the C_P_ Thiolate nucleophilicity increased by Arg and Thr hydrogen bonding networks **2.** Hydroperoxide (Hpx) trapped by Arg which makes it to target in active site **3.** Shift of Arg, Thr H bonds from C_P_ to Hpx stabilizes the Hpx & increase the thiolate reactivity, enabling SN2 reaction **4.** C_P_ is oxidised to C_P_–SOH & Arg polar gp stabilizes the Leaving gp **5**. SOH triggers structural rearrangement & allows condensation among C_P_ -C_R_ **6.** Disulfide bond formation & release of water as a product.

## 4. Conclusion

In this study, we have elucidated, for the first time, the critical role of the N-terminal β-flap region in bacterial thiol peroxidases (Tpxs) as an active site distant regulator of its enzymatic activity. Through a combination of high-resolution X-ray crystallography, biochemical characterization, and mutational analysis, we demonstrate that this structurally conserved motif stabilizes the catalytic triad (Cys60, Thr57, Arg128) by maintaining the spatial orientation of key residues essential for substrate binding and catalysis. Deletion of the β-flap (NΔ15-SaTpx) disrupts this architecture, leading to a distorted active site geometry, impaired redox activity, and compromised oxidative stress resilience. Comparative structural analyses with mammalian orthologs (e.g., human Prx5) further reveal that the N terminal β-flap is a bacterial-specific feature, absent in eukaryotic counterparts, suggesting its evolutionary specialization in prokaryotic redox defense systems. These findings not only advance our mechanistic understanding of bacterial peroxiredoxin function but also identify the β-flap as a potential target for structure-guided inhibitor design, offering a pathway to disrupt pathogen-specific antioxidant mechanisms without affecting host enzymes. This work underscores the therapeutic potential of targeting novel regulatory sites in bacterial redox enzymes to combat infections caused by multidrug-resistant pathogens.

## Credit, authorship contribution statement

**Manjari Shukla:** investigation, data curation, methodology, analysis, writing original draft. **Sushobhan Maji:** data curation. **Amit Kumar Das:** data curation, conceptualization**. Amit Mishra:** data curation, analysis. **Sudipta Bhattacharyya:** writing, editing, review, methodology, validation, supervision, conceptualization, resources.

## Acknowledgements

The authors thank X-ray facility at Raja Ramanna Centre for Advanced Technology (RRCAT), Indore, India for providing access to their instrumentation and technical support. The research work is supported by SEED Grant (I/SEED/SUB/20200005) sponsored by IIT Jodhpur and Startup Research Grant (S/SERB/SUB/20200055) funded by Science & Engineering Research Board (SERB), Department of Science & Technology, India.

