## Supplementary data for "Shaping the active site from a distance: Unravelling the regulatory role of the conserved N-terminal β-flap of thiol peroxidases through structure-function characterization of the Staphylococcal ortholog"

### 2. Material and Method

#### 2.1 Reagents, fine chemicals and enzymes

The Luria broth, Tris, Imidazole, NaCl, Coomassie Brilliant Blue R250, Bromophenol Blue, β-mercaptoethanol, Glacial acetic acid, Ethanol, Methanol, Sodium dodecyl sulphate, Bis-acrylamide Acrylamide, Ammonium persulfate, TEMED, DTNB, L-cystine, NADPH, Tris base, NaCl, and Imidazole purchased from SRL (Mumbai, India). EDTA, Agarose, Glycerol, Calcium chloride, IPTG, dithiothreitol, Kanamycin, Ampicillin, dNTPs, 100bp DNA ladder and 24 well plates purchased from HiMedia (Mumbai, India). DNAse free water, enzymes such as, *Taq* DNA polymerase, *Bam*HI, *Sal*I, T4 DNA ligase procured from NEB (New England Biolabs, United States). *Pfu* DNA polymerase, Protein molecular weight marker supplied by Promega (United States). Kits like PCR purification kit, Plasmid miniprep kit, Vectors like pET28a, pQE30 and Cells like BL21(DE3), M15 and SG13009 purchased from Qiagen (Germany). Column matrix like Ni-Sepharose high performance matrix, Superdex 75, Superdex 200 matrix procured from Cytiva (United States). Tert-butyl hydroperoxide and cumene hydroperoxide purchased from Sigma Aldrich (USA) while hydrogen peroxide was purchased from Qualigens (Mumbai India) and Vivaspin 20 concentrator (10kDa MWCO) for protein concentration was procured from GE Healthcare. The sparse matrix screening solutions (Crystal Screen I-HR2--110, Crystal Screen II-HR2-112, and Index Screen-HR2-144), 96 well Linbro plates, Dow Corning grease and Rain-X have been procured from Hampton Research USA, and round coverslips from BlueStar India. The other analytical grade fine chemicals for fine screening of the initial crystallization conditions have been procured from Sigma-Aldrich USA.

#### 2.2. Cloning of Staphylococcal Thiol Peroxidase (SaTpx), Thiol Peroxidase N- terminal deletion mutant (NΔ15-SaTpx), Thioredoxin reductase1 and Thioredoxin reductase (SaTR)

Genomic DNA of *S. aureus* RN4220 has been used as the template for amplification of SaTpx (SAOUHSC 01822), NΔ15-SaTpx, SaTrx1 (SAOUHSC 01100) and SaTR (SAOUHSC 00785) open reading frame by using the below set of primers and restriction enzymes recognition sites.

**Table S1: Primer designed for target gene cloning**

| **Target Gene** | **Forward primer with BamHI restriction site (in bold letters)**<br>**Reverse primer with SalI restriction site (in bold letters)** |
|---|---|
| SaTpx | 5’-ATATAT**GGATCC**ATGACTGAAATAACATTCAA-3’<br>5’-ATATAT**GTCGAC**TTAAATATTTTTGTATGCAG-3’ |
| NΔ15-SaTpx | 5’ ATATAT**GGATCC**TCAAAGGTGGACCAA-3’<br>5’-ATATAT**GTCGAC**TTAAATATTTTTGTATGCAG-3’ |
| SaTrx1 | 5’-ATATAT**GGATCC**ATGGCAATCGTTAAAGTA-3’<br>5’-ATATAT**GTCGAC**TTATAAATGTTTATCTAA-3’ |
| SaTR | 5’-ATATAT**GGATCC**ATGACTGAAATAGATTTTGA-3’<br>5’-ATATAT**GTCGAC**TTAAGCTTGATCGTTTAAAT-3’ |

### 3. Results and Discussion

SaTpx (UniProt Id: Q2FXL3), NΔ15-SaTpx, SaTrx1 (UniProt Id_Q2FZD2) and SaTR (UniProt Id_Q2G041)

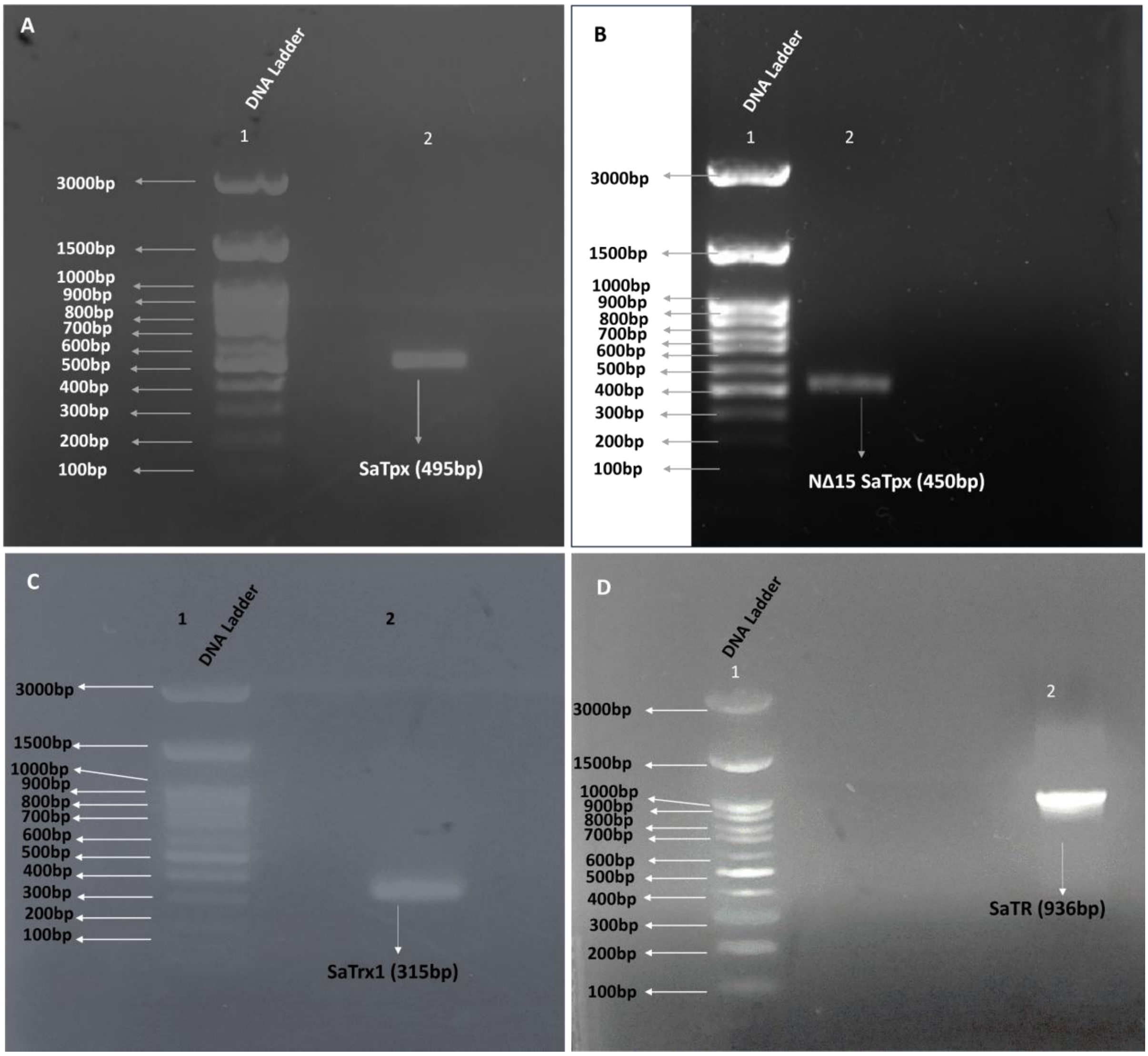


**Figure S1: Agarose gel electrophoresis of amplified gene of interest. 1%(W/V) agarose gel electrophoresis profile showing the PCR amplicons of the targetd proteins’ ORF.** A) SaTpx (495bp) B) NΔ15-SaTpx (450bp) C) SaTrx1(315bp) and D) SaTR (936bp)

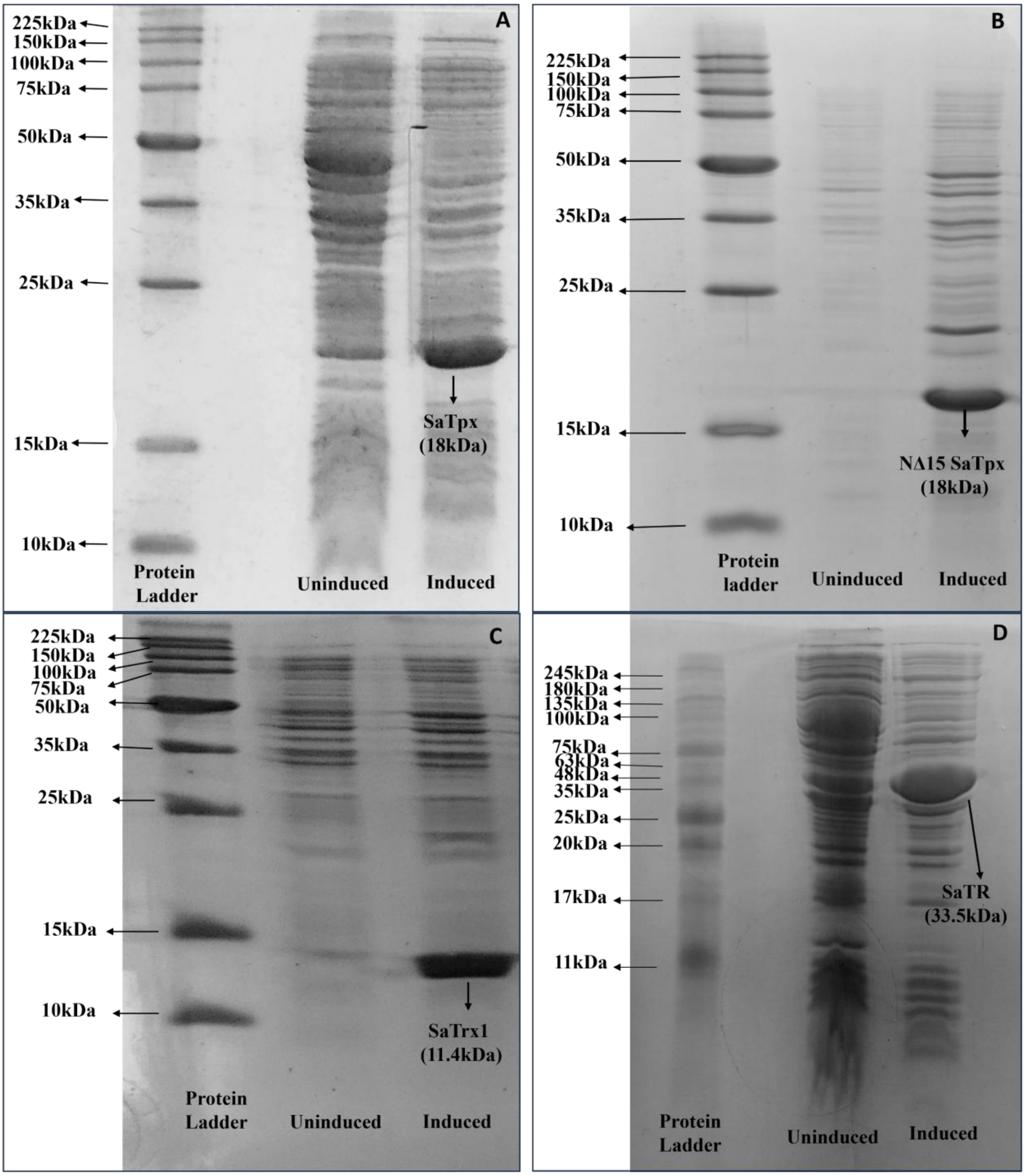


**Figure S2: Overexpression of proteins under IPTG induction. SDS PAGE profile showing overexpression of target proteins at under optimized IPTG induction**: The diagram shows the uninduced and induced protein bands in 12-15% SDS PAGE gels. A) SaTpx, B) NΔ15-SaTpx C) SaTrx1 and D) SaTR

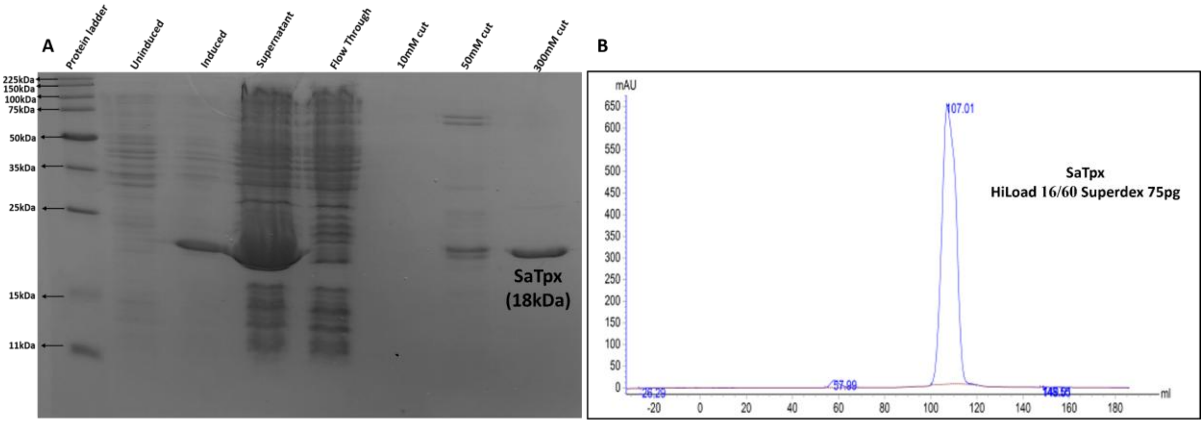


**Figure S3: Purification profile of recombinant Staphylococcal Peroxiredoxin enzyme (SaTpx). S3A:** 15% SDS PAGE profile showing purification of SaTpx through IMAC. The protein was eluted from the IMAC column using different step-gradients of imidazole. **S3B:** UV280nm trace showing sharp single Gaussian peak of gel filtration chromatography (GFC) purified SaTpx.

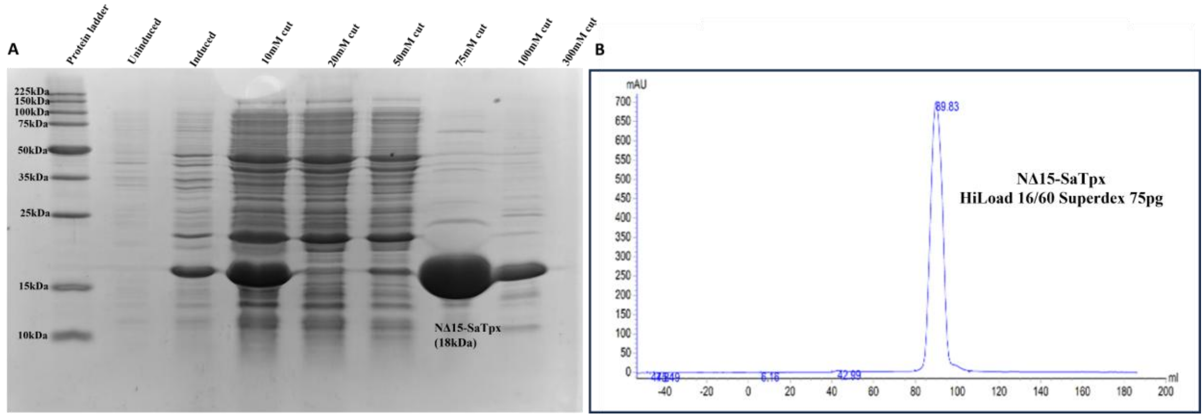


**Figure S4: Purification profile of recombinant N terminal deletion mutant of Staphylococcal Peroxiredoxin enzyme (NΔ15-SaTpx). S4A:** 15% SDS PAGE profile showing purification of NΔ15-SaTpx through IMAC. The protein was eluted from the IMAC column using different step-gradients of imidazole. **S4B:** UV280nm trace showing sharp single Gaussian peak of gel filtration chromatography (GFC) purified NΔ15-SaTpx.

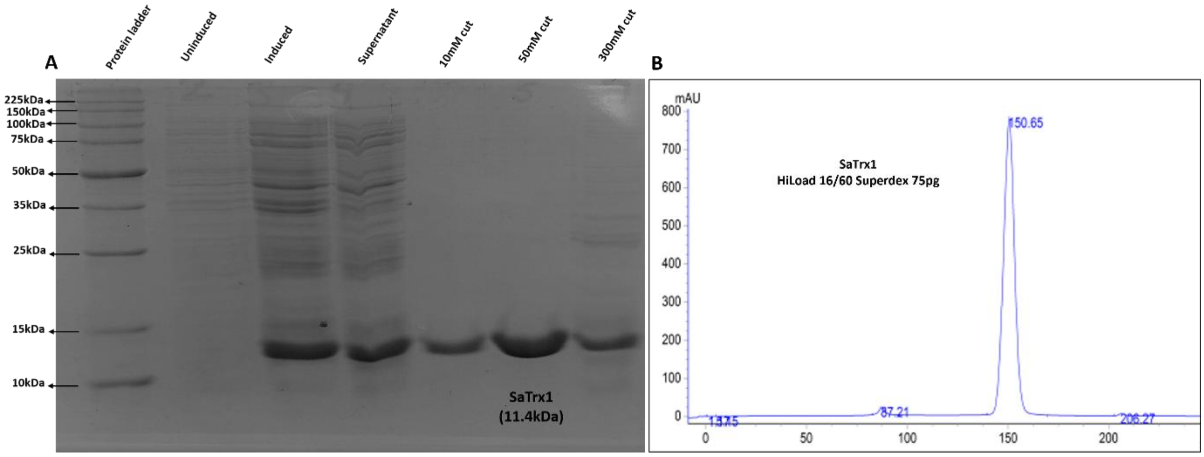


**Figure S5: Purification profile of recombinant Staphylococcal Thioredoxin 1 enzyme (SaTrx1). S5A:** 12% SDS PAGE profile showing purification of SaTrx1 through IMAC. The protein was eluted from the IMAC column using different step-gradients of imidazole. **S5B:**

UV280nm trace showing sharp single Gaussian peak of gel filtration chromatography (GFC) purified SaTrx1.

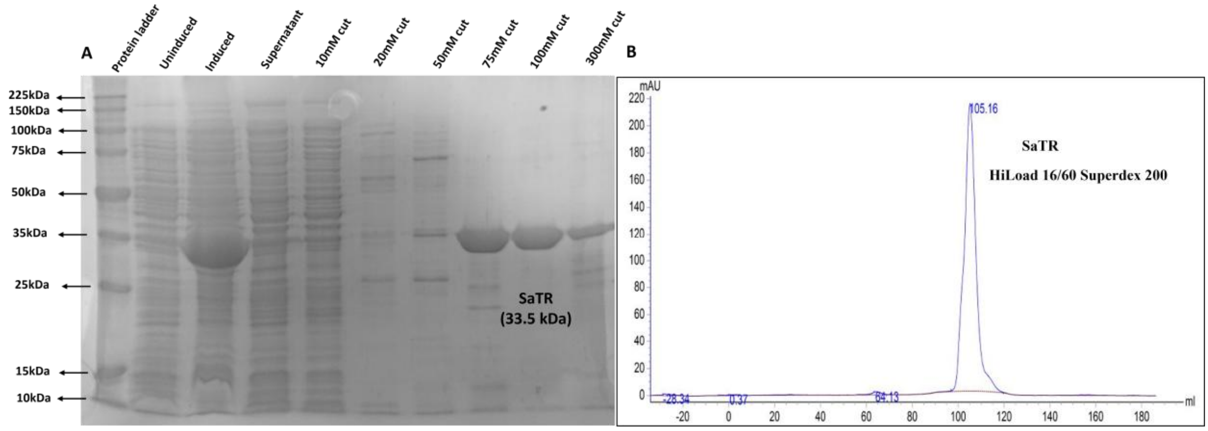


**Figure S6: Purification profile of recombinant Staphylococcal Thioredoxin Reductase enzyme (SaTR). S6A:** 12% SDS PAGE profile showing purification of SaTR through IMAC. The protein was eluted from the IMAC column using different step-gradients of imidazole. **S6B:** UV280nm trace showing sharp single Gaussian peak of gel filtration chromatography (GFC) purified SaTR.

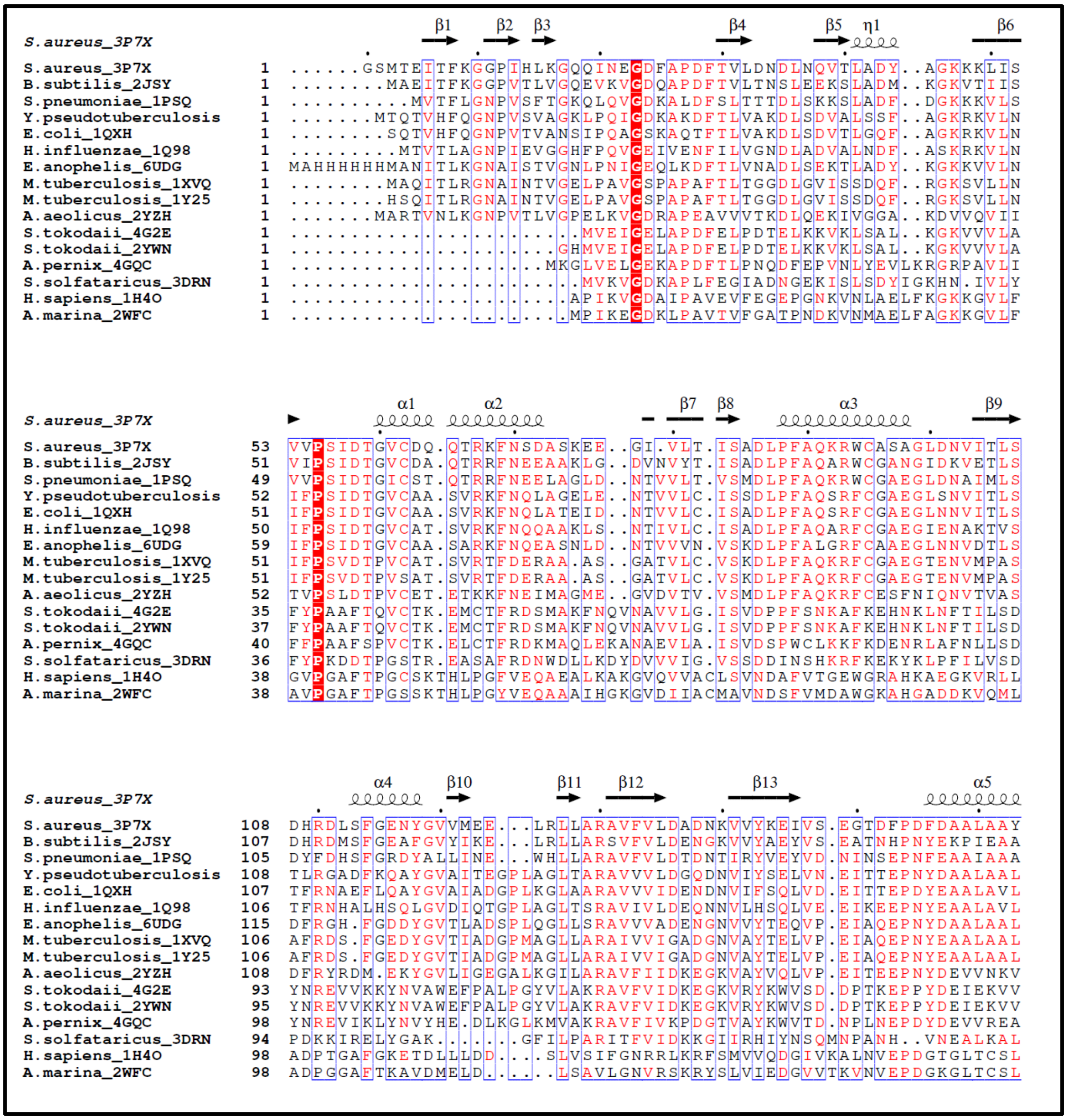


**Figure S1b**_Figure 1b (extended form): Representative PDB structures of atypical 2 cysteine peroxiredoxin from subfamilies of peroxiredoxins (Tpx, BCP, PrxQ, Prx5). The N terminal β flap is present in prokaryotes only.

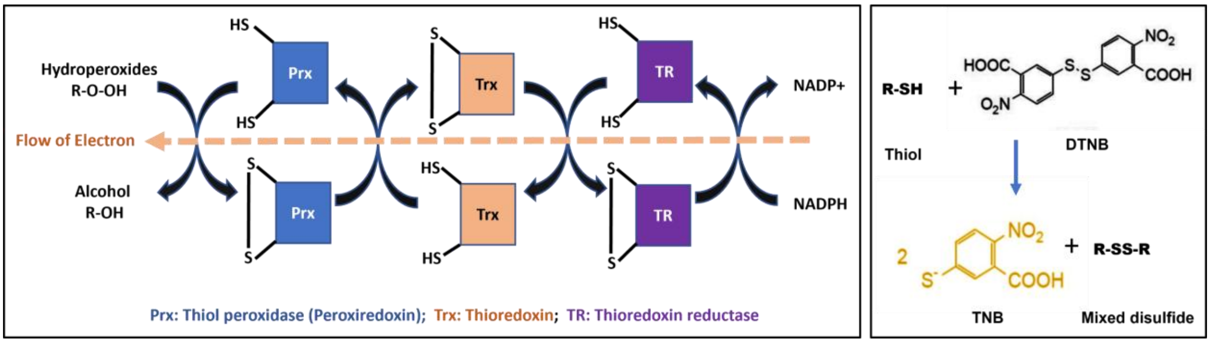


**Figure S2e (inset of Figure 2a, 2b):** Mechanism of the Thioredoxin-Dependent Peroxiredoxin Catalytic Cycle and Principle of DTNB-based Thiol Detection.

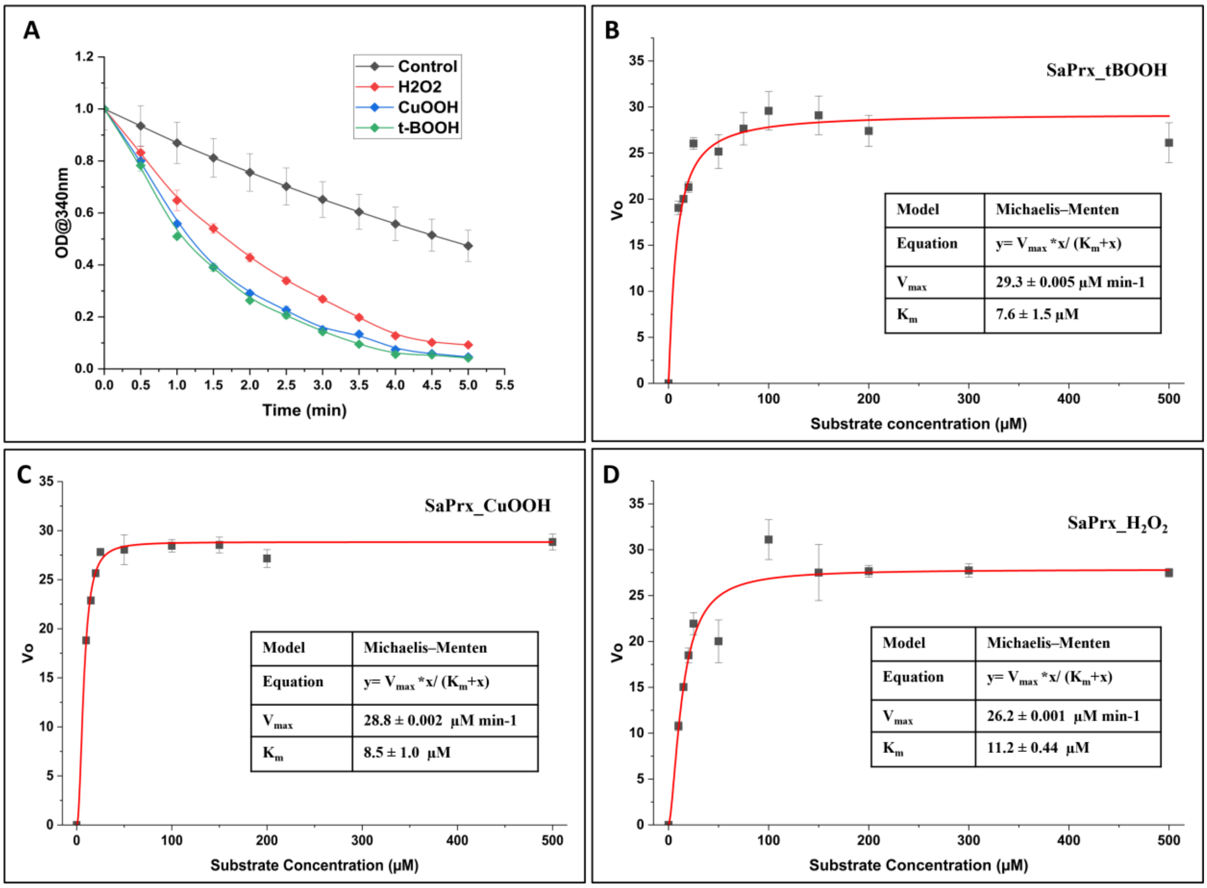


**Figure S7: The enzyme kinetic parameters of the purified SaTpx Enzyme. S7A:** Time-dependent NADPH oxidation kinetics with various peroxide substrates. Spectrophotometric monitoring at 340 nm demonstrates differential rates of NADPH consumption in a coupled enzyme assay system. Tert-Butyl hydroperoxide (t-BOOH) exhibited the most rapid NADPH depletion. Error bars represent standard deviation (n=3). **Figure S7B-D:** Michaelis-Menten kinetic analysis of SaTpx with three distinct peroxide substrates. Initial velocity ($V_0$) as a function of substrate concentration (0-500 µM) was fitted to the Michaelis-Menten equation ($V=V_{max}\cdot[S]/(K_m+[S])$). The enzyme demonstrated differential substrate preferences as evidenced by their respective kinetic parameters.

**Table S2: Comparative enzyme kinetic parameters of SaPrx against different hydroperoxide substrates**

| **Parameter ⟶ Enzymes ↓** | **$K_m$ (M)** | **$V_{max}$ (M $sec^{-1}$)** | **$K_{cat}$ = $V_{max}/[E]$ ($sec^{-1}$)** | **$K_{cat}/K_m$ ($M^{-1}$ $sec^{-1}$)** |
|---|---|---|---|---|
| **SaPrx_tBOOH** | 7.6 x $10^{-6}$ ±1.5 | 4.88 x$10^{-7}$± 0.005 | 9.7 x $10^{-2}$ | 12.8 X $10^{3}$ |
| **SaPrx_CuOOH** | 8.5 x $10^{-6}$ ± 1.03 | 4.8 x$10^{-7}$ ± 0.002 | 9.6 x $10^{-2}$ | 11.3 × $10^{3}$ |
| **SaPrx_ $H_2O_2$** | 11.2 x $10^{-6}$ ± 0.4 | 4.37 x$10^{-7}$± 0.001 | 8.7 x $10^{-2}$ | 7.8 × $10^{3}$ |

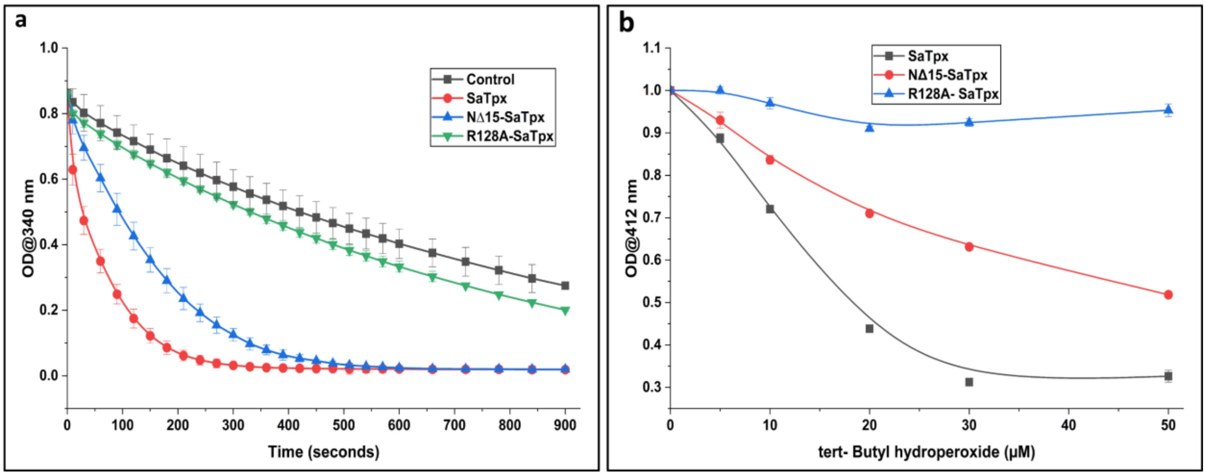


**Figure S8: Comparative enzyme catalysis and Kinetic Characterization of Wild-Type and Mutant *Staphylococcus aureus* Peroxiredoxin (SaTpx and NΔ15-SaTpx). S8a: Time-course analysis of NADPH consumption in a coupled peroxidase assay**. Absorbance at 340 nm, decreases over time for SaTpx (red), NΔ15-SaTpx (blue), R128A-SaTpx (green) and the control (black-no peroxidase), demonstrating enzyme-mediated NADPH oxidation. **S8b: Substrate-dependent oxidation of thiol groups in SaTpx and NΔ15-SaTpx.** The plot shows the decrease in absorbance at 412 nm with increasing tert-butyl hydroperoxide (t-BOOH) concentration, indicating oxidation of free thiols. SaTpx exhibits a steeper decline compared to NΔ15-SaTpx, reflecting higher reactivity towards t-BOOH whereas R128A mutant shows no activity.

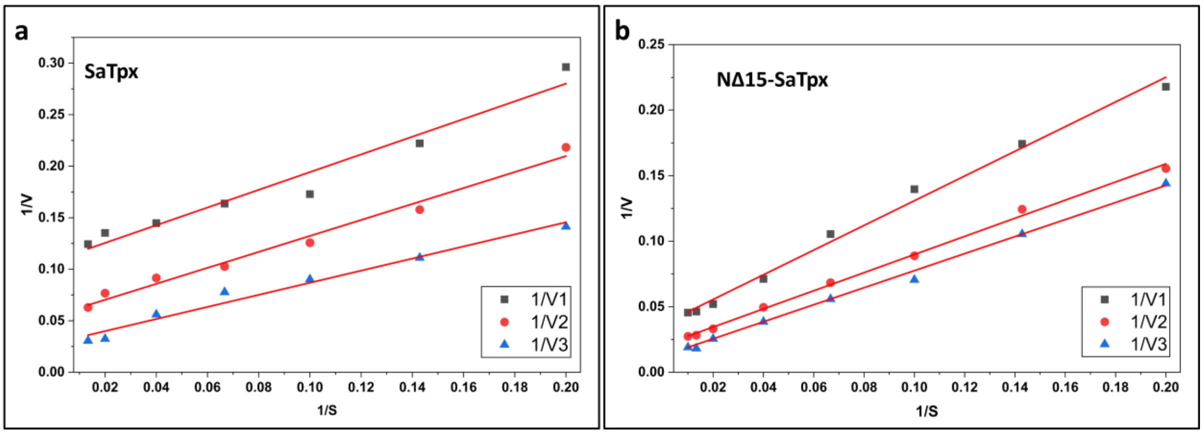


**Figure S9:** Double reciprocal plot (1/S vs 1/V): a: SaTpx double reciprocal plot at three Trx1 concentrations b: NΔ15-SaTpx double reciprocal plot at three Trx1 concentrations

From the above double reciprocal plot, we have calculated the true kinetic constants ($K_m$ and $V_m$, $K_{cat}$ and $K_{cat}/K_m$) of SaTpx and NΔ15-SaTpx.

Table S3: True enzyme kinetic constants for SaTpx and NΔ15-SaTpx

| **Parameter →** **Enzymes ↓** | **$K_m$ (Peroxide) (M)** | **$V_{max}$ ($M\ sec^{-1}$)** | **$K_{cat}$ = $V_{max}/[E]$ ($sec^{-1}$)** | **$K_{cat}/K_m$ ($M^{-1}\ sec^{-1}$)** |
|---|---|---|---|---|
| SaTpx | $20 \times 10^{-6}$ | $11.1 \times 10^{-6}$ | 2.22 | $11.1 \times 10^{4}$ |
| NΔ15-SaTpx | $59 \times 10^{-6}$ | $3.3 \times 10^{-6}$ | 0.65 | $1 \times 10^{4}$ |

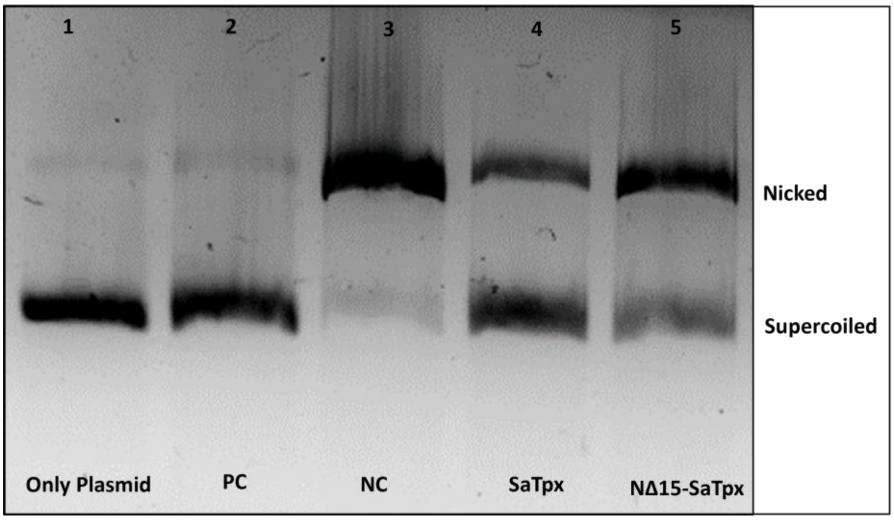


**Figure S10: ROS induced dsDNA protection activity of peroxiredoxin proteins.** Metal-catalysed ROS induced Plasmid DNA nicking and its protection by SaTpx, NΔ15-SaTpx was visualized by 1% agarose gel electrophoresis demonstrating the antioxidant capacity of purified peroxiredoxins. Lane 1: untreated plasmid DNA (supercoiled control); Lane 2: positive control where EDTA added before the start of the reaction to quench the metal ions (PC); Lane 3: negative control having no SaTpx enzymes; Lane 4: SaTpx-treated sample; Lane 5: NΔ15-SaTpx -treated sample. The migration positions of nicked (oxidatively damaged) and supercoiled (protected) DNA conformations are indicated on the right.

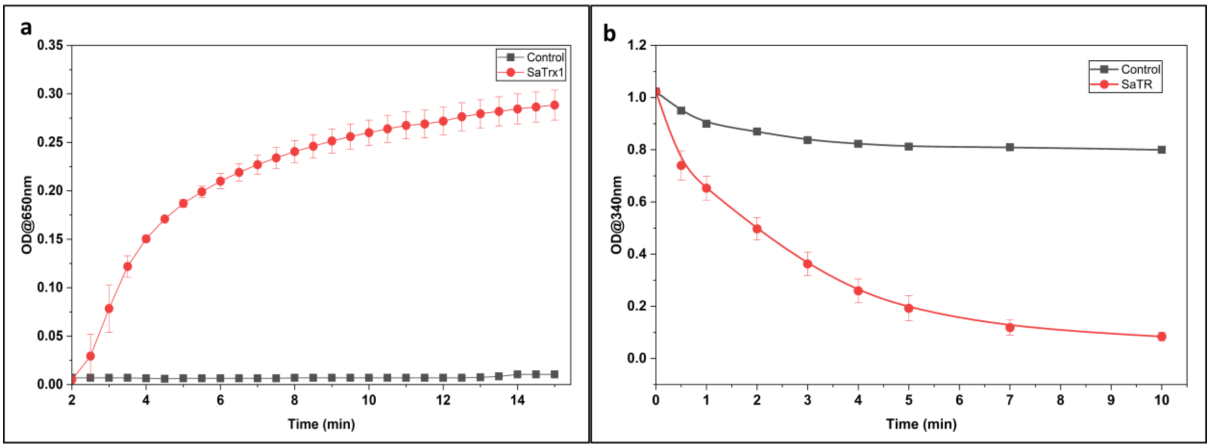


**Figure S11a:** Turbidimetric assay for Insulin reduction by Trx1 were comparing SaTrx1 (red) and control (black), the progressive increase in absorbance reflects the time dependent precipitation of reduced insulin indicating efficient reduction of Insulin by Trx1 thus suggesting activity of Trx1. **Figure S11b:** NADPH oxidation assay for SaTR: The red circles show decreasing OD at 340nm over the time, indicating rapid consumption of NADPH. By contrast control (black) maintains OD340 over time, confirms the NADPH oxidation depends on active SaTR.

Together these plots suggest the functional activity of SaTrx1 in catalysing disulfide bond reduction and SaTR in catalysing NADPH dependent thioredoxin reduction.

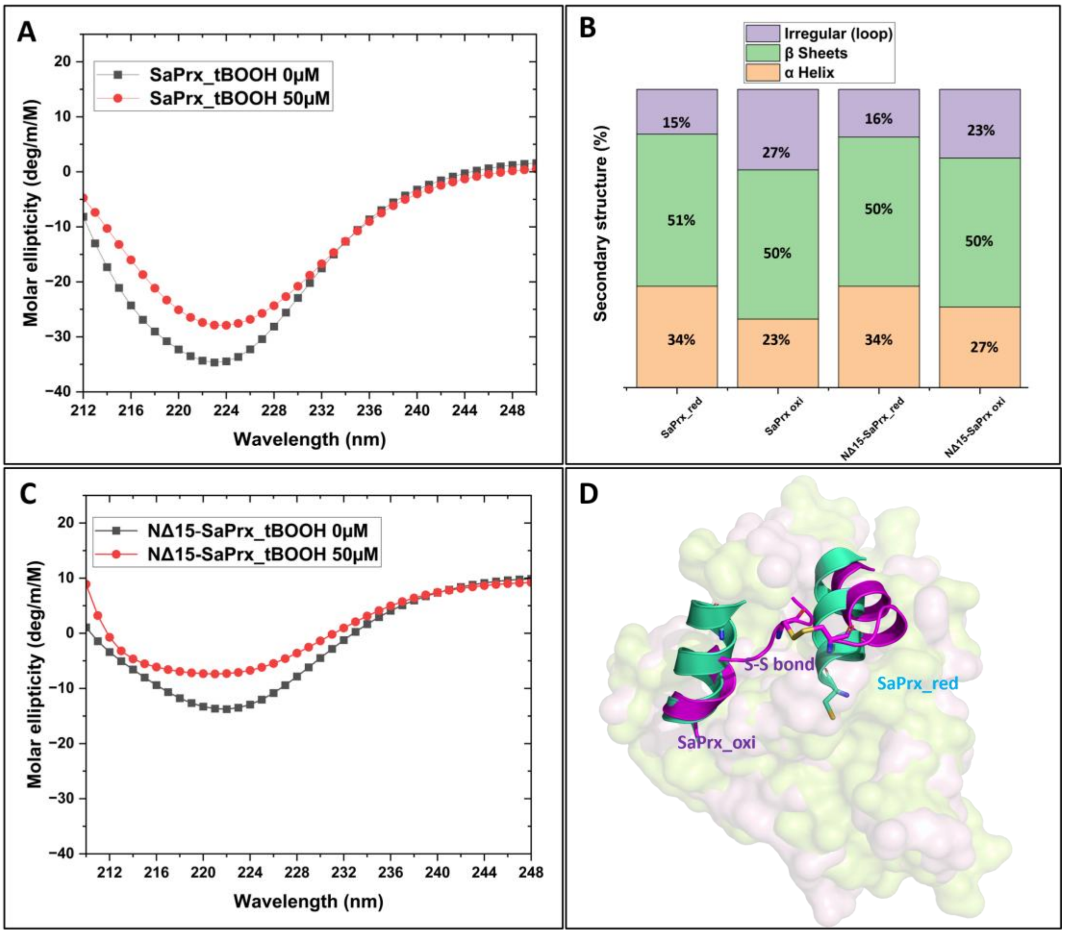


**Figure S12: Oxidation-induced secondary structural changes of SaTpx and NΔ15-SaTpx enzymes. Figure S12A: Far-UV circular dichroism spectroscopic analysis of SaTpx in reduced state (0µM tBOOH, black squares) and oxidized state (50µM tert-butyl hydroperoxide, red circles).** The spectra exhibit characteristic minima at approximately 222 nm, with reduced SaTpx displaying pronounced negative ellipticity (-35 $deg\cdot cm^2\cdot dmol^{-1}$) compared to its oxidized counterpart (-28 $deg\cdot cm^2\cdot dmol^{-1}$), indicating substantial α-helical content alteration upon oxidation. **Figure S12C: Far-UV circular dichroism analysis of N-terminal loop-deleted mutant NΔ15-SaTpx in reduced state (black squares) and oxidized state (red circles).** The mutant also exhibits the spectral changes upon oxidation like wild-type protein, with reduction in negative ellipticity at the 222 nm minimum (-13 $deg\cdot cm^2\cdot dmol^{-1}$ to -8 $deg\cdot cm^2\cdot dmol^{-1}$), suggesting conformational responsiveness to oxidative modification. **Figure S12B: Quantitative secondary structure composition analysis of SaTpx and NΔ15-SaTpx in reduced and oxidized states**. Deconvolution of CD spectra reveals significant alterations in structural elements upon oxidation: wild-type SaTpx shows reduction in α-helical content (34% to 23%) accompanied by increased irregular structures i.e loop (15% to 27%), while NΔ15-SaTpx also demonstrates α-helical loss (34% to 27%) with substantial increase in irregular structures (16% to 23%). β-sheet content remains relatively stable in both proteins, suggesting preservation of core structural elements during redox transitions. **Figure S12D: Homology model based Structural superposition of reduced (teal) and oxidized (magenta) SaTpx.** The cartoon representation illustrates the conformational rearrangement induced by oxidation, highlighting the formation of the disulfide bond (stick representation) between Cys60 and Cys93. This redox-dependent structural transition represents the catalytic mechanism of atypical 2-Cys peroxiredoxins and correlates with the secondary structural changes observed spectroscopically in panels A-C.

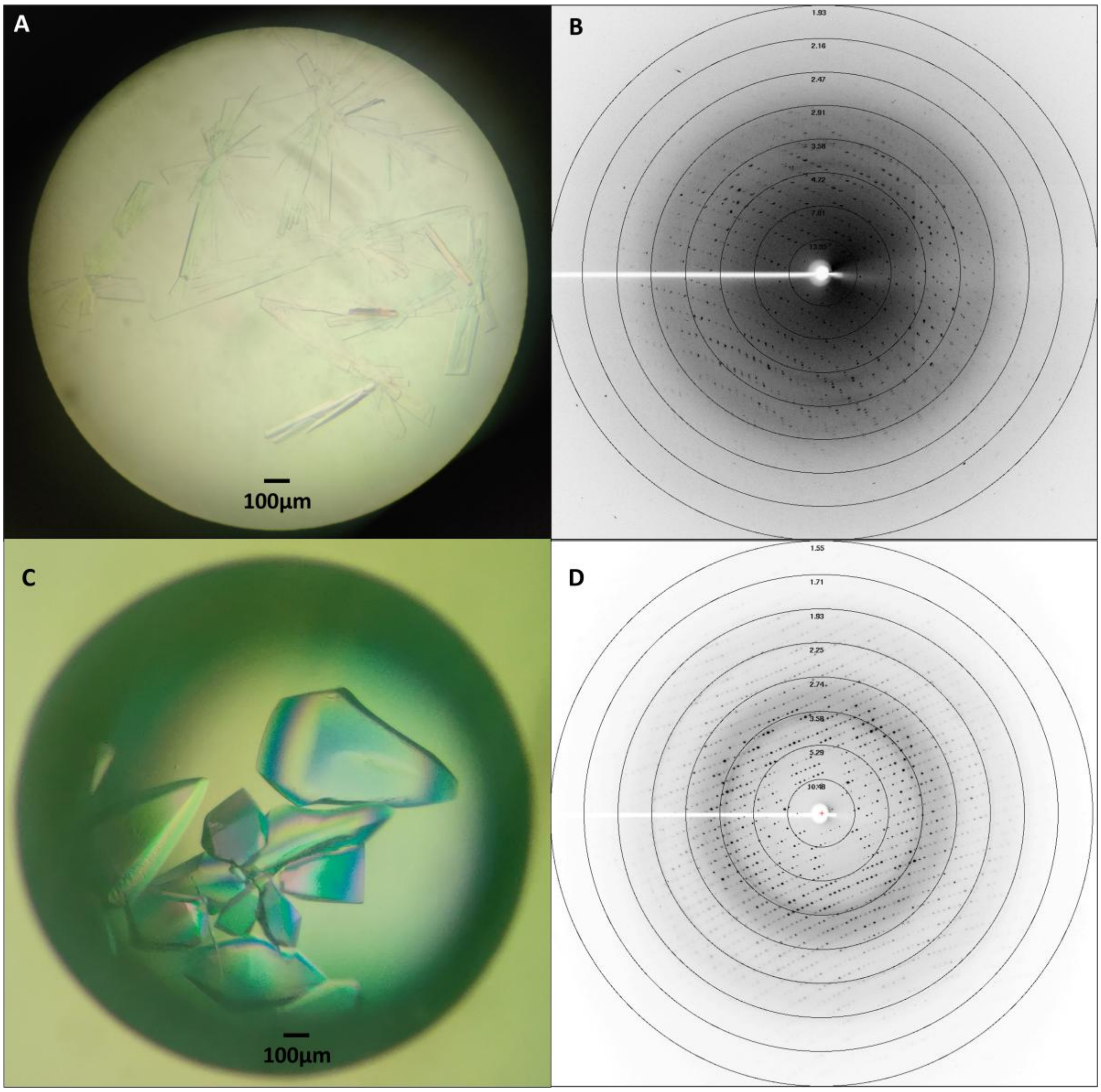


**Figure S13: Crystallization and Single crystal X-ray diffraction of the target proteins. Figure S13A:** Diffraction quality single crystals of SaTpx produced via vapor diffusion hanging drop method. **Figure S13B: X**-ray diffraction pattern of SaTpx crystals, shows well-defined spots extending to a resolution of ~ 1.8 Å. **Figure S13C:** Diamond-shaped, well-formed crystals of NΔ15-SaTpx grown at room temperature via the vapor diffusion hanging drop technique. **Figure S13D:** X-ray diffraction spots of NΔ15-SaTpx crystals soaked with the substrate mimic OtBH, exhibiting diffraction spots upto 1.55 Å resolution, suitable for high-resolution structural determination.

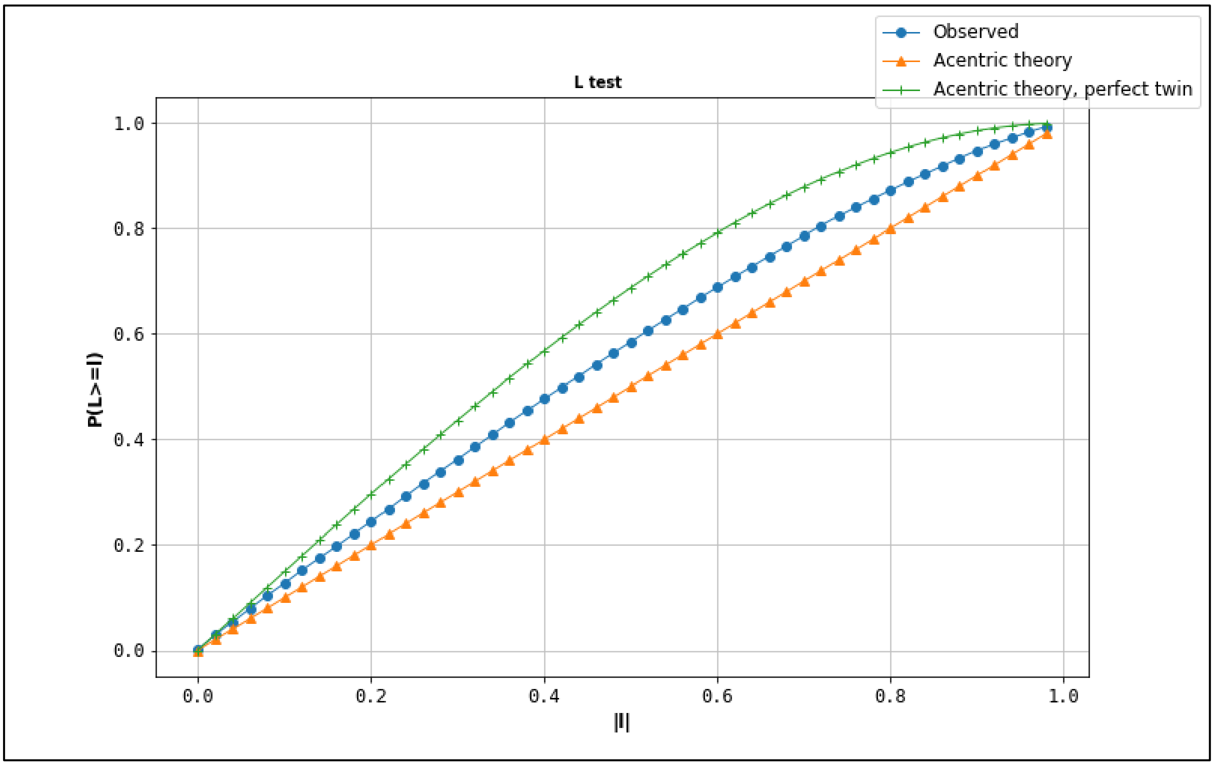


**Figure S14: L-test analysis of OtBH bound NΔ15-SaTpx diffraction data**: L-Test analysis for twinning detection in diffraction data by Phenix Xtriage gives cumulative intensity plot which indicates the partial twinning as the observed data (blue line) is deviated from the untwined theoretical distribution (orange line).

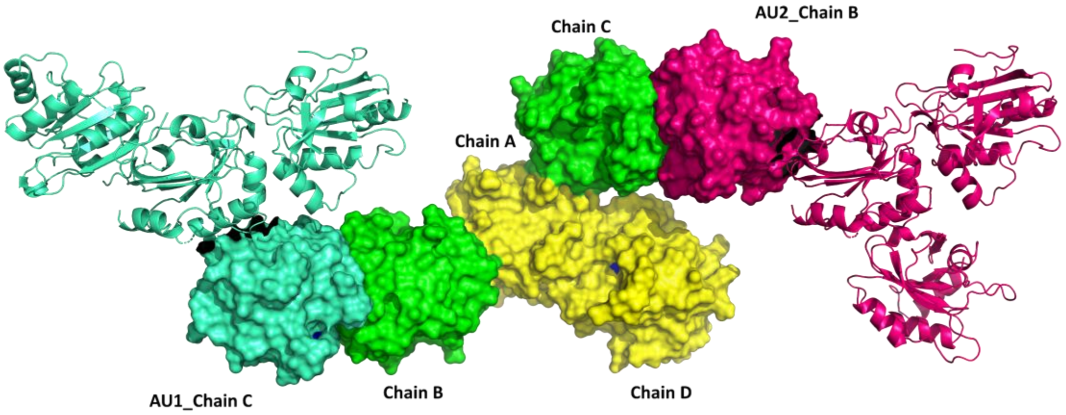


**Figure S15:** Two additional monomers also make functional dimeric architecture with the neighbouring monomers of the adjacent unit cells. The monomers of one unit forming functional dimeric interface (chain A/D_yellow surface) and other two monomers (chain B& C, green surface) forming functional dimeric interface with adjacent unit cells (magenta (adjacent unit 2) and cyan (adjacent unit 1) surface representation).

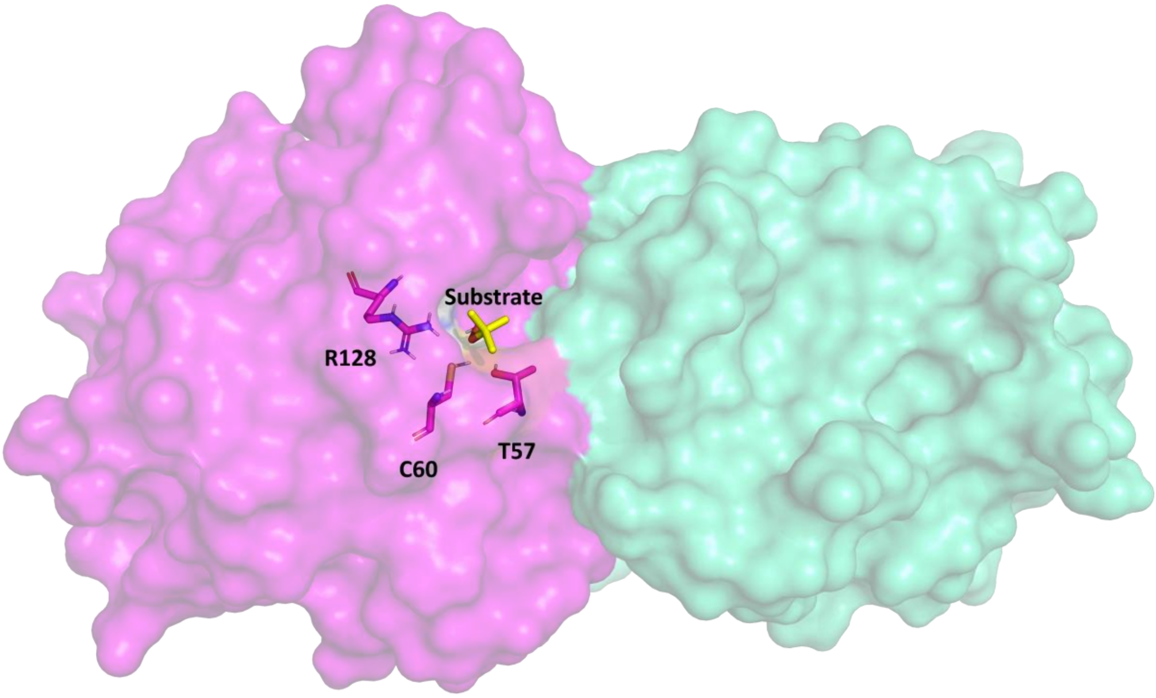


**Figure S16:** The close proximity of the active site catalytic triad amino acid residues (represented in stick purple colour) to the SaTpx dimeric (Chain A_purple & Chain B_cyan surface representation) interface demonstrates the invincibility of the dimeric interface for the formation of the active site catalytic cleft (cleft binding substrate, yellow stick) of SaTpx.

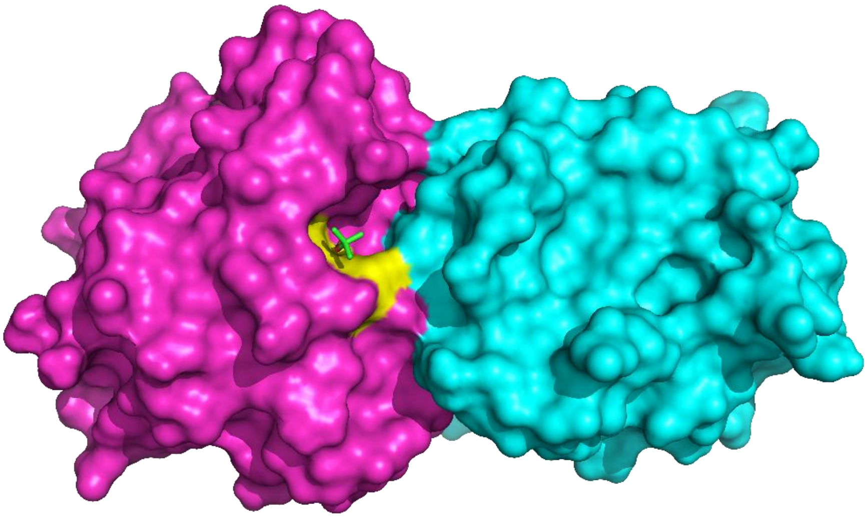

**Figure S17:** Elongated active site architecture (L shaped) of bacterial Tpxs docked with tert-butyl hydroperoxide substrate represented as cyan stick.

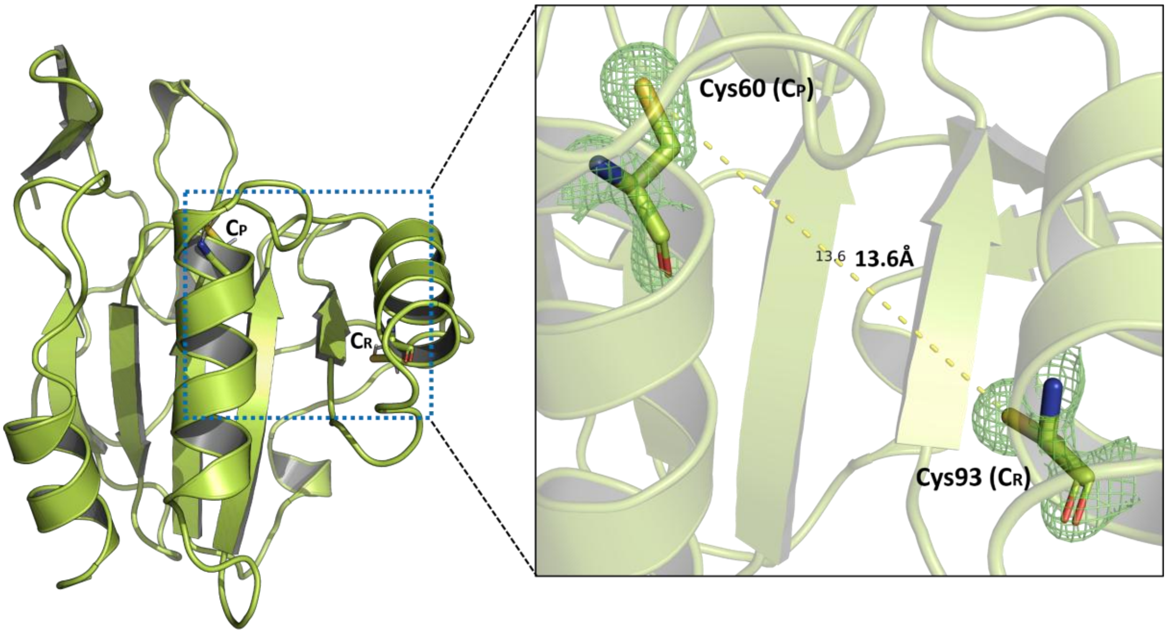


**Figure S18:** In SaTpx crystal structures these two cysteines ($C_P$ and $C_R$ are C60 and C93 respectively, represented with stick) are located on α2 and α3 helices respectively, with a Cα distance of ~13Å with respect to each other.

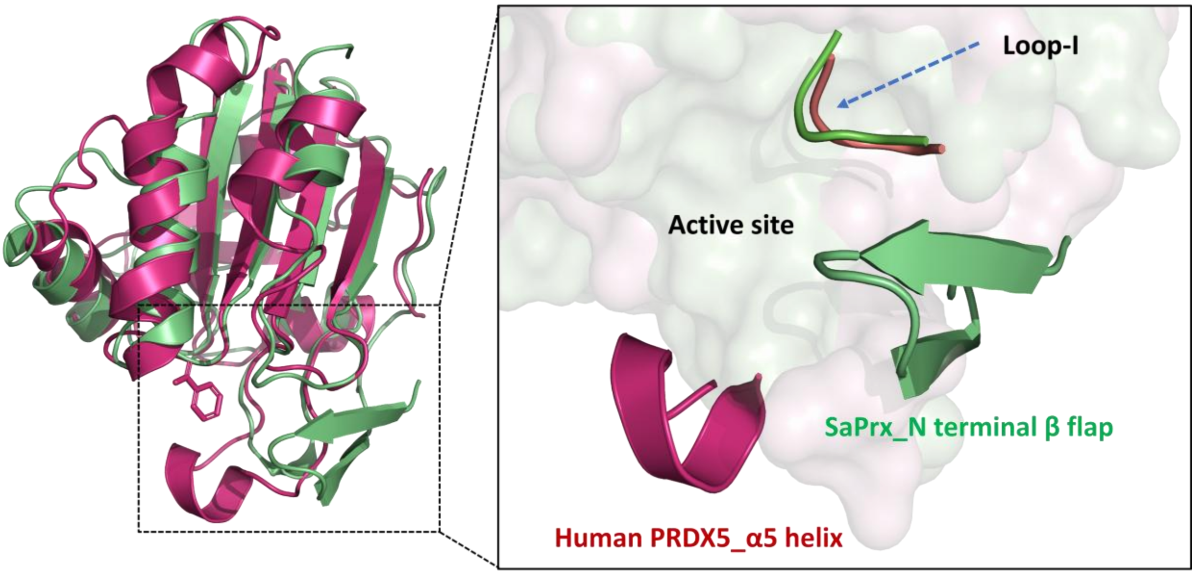


**Figure S19: Structural variations in SaTpx and mammalian atypical 2 cysteine peroxiredoxin:** Structural alignment of mutant SaTpx (green) with human PRDX5 (pink) reveals that PRDX5 retains its Loop-I in close proximity to the catalytic triad (as observed in WT-SaTpx) within the active site. Notably, in PRDX5, the loop connecting the α4 helix and β6 strand extends and folds into a distinct short α5 helix (red), which is proposed to function analogously to the N-terminal β-flap of SaTpx, contributing to active site architecture and stabilization.

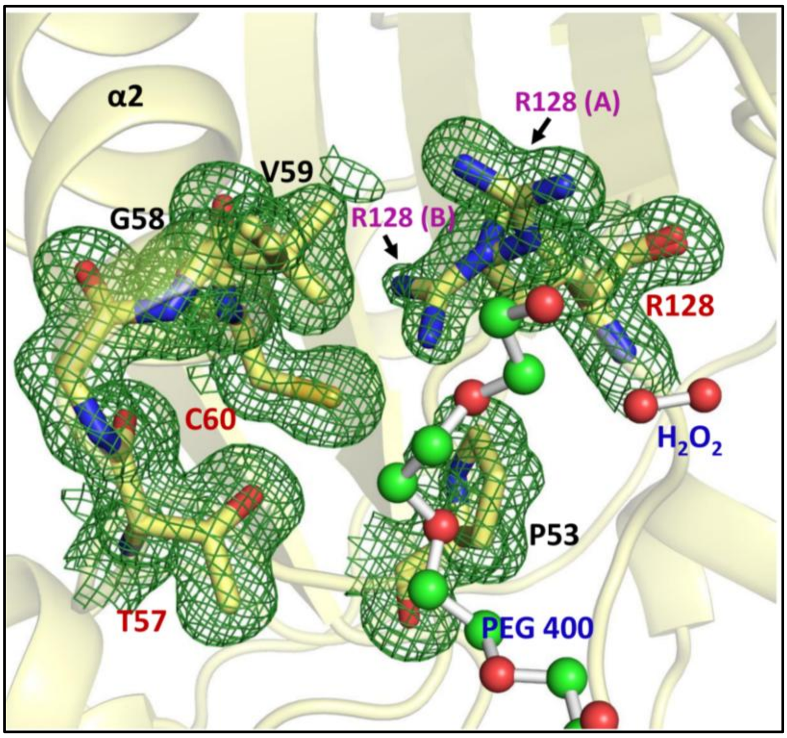


**Figure S20: Electron density (2Fo−Fc map contoured at 1σ) around the active-site catalytic triad of** NΔ15-SaTpx mutant solved at 1.25Å resolution: R128 shows dual conformations of its side chain guanidinium group ([(H2N)(HN)CN(H)-): one oriented toward C60 thiol group (-SH) forming a hydrogen bond (R128 B), and in the alternative conformation R128 guanidinium group is flipped away, abolishing this interaction.
